# The Chinese chestnut genome: a reference for species restoration

**DOI:** 10.1101/615047

**Authors:** Margaret Staton, Charles Addo-Quaye, Nathaniel Cannon, Yongshuai Sun, Tetyana Zhebentyayeva, Matthew Huff, Shenghua Fan, Emily Bellis, Nurul Islam-Faridi, Jiali Yu, Nathan Henry, Anna Conrad, Daniela I. Drautz-Moses, Xingfu Zhu, Zhiqiang Lu, Rooksana E. Noorai, Stephen Ficklin, Chris Saski, Mihir Mandal, Tyler K Wagner, Nicole Zembower, Catherine Bodénès, Jason Holliday, Jared Westbrook, Jesse Lasky, Laura Georgi, Fred V Hebard, C. Dana Nelson, Stephan C Schuster, Albert G Abbott, JE Carlson

## Abstract

Forest tree species are increasingly subject to severe mortalities from exotic pests, diseases, and invasive organisms, accelerated by climate change. Forest health issues are threatening multiple species and ecosystem sustainability globally. While sources of resistance may be available in related species, or among surviving trees, introgression of resistance genes into threatened tree species in reasonable time frames requires genome-wide breeding tools. Asian species of chestnut (*Castanea* spp.) are being employed as donors of disease resistance genes to restore native chestnut species in North America and Europe. To aid in the restoration of threatened chestnut species, we present the assembly of a reference genome with chromosome-scale sequences for Chinese chestnut (*C. mollissima*), the disease-resistance donor for American chestnut restoration. We also demonstrate the value of the genome as a platform for research and species restoration, including new insights into the evolution of blight resistance in Asian chestnut species, the locations in the genome of ecologically important signatures of selection differentiating American chestnut from Chinese chestnut, the identification of candidate genes for disease resistance, and preliminary comparisons of genome organization with related species.

## INTRODUCTION

Genome resources hold much promise for, and may provide the key to, the restoration of forest tree species which have been, or are in the process of being, extirpated from their natural habitats by environmental threats imposed by exotic pests and diseases, invasive organisms, and climate change. These threats, generally referred to as Forest Health issues, are responsible for severe mortalities in numerous forest tree species. The extirpation of American chestnut from its natural range by the invasive Asian pathogens *C. parasitica* and *P. cinnamomi* in the first half of the 20^th^ century was recognized as the greatest environmental disaster of the time (1). To address these and other environmental challenges (e.g. rapid climate change), we have assembled a Chinese chestnut reference genome to aid in the transfer of chestnut blight resistance loci from for Chinese chestnut to American chestnut through introgression (back-cross breeding) and potential biotechnology approaches, and to serve as a model for other genome-assisted species restoration efforts in long-lived, undomesticated plant species.

Chestnuts (*Castanea* species) are members of the Fagaceae (Order Fagales), whose members include many important forest tree species worldwide. The Fagaceae is comprised of eight genera containing 1,105 accepted species names (The Plant List, www.theplantlist.org). These species are not only important to human industry (timber, pulp wood, furniture and others) but are also dominant species in many of our forested ecosystems, providing food, and shelter for wildlife as well as other important ecosystem services. Within *Castanea* the seven recognized species - *C. crenata*, *C. dentata*, *C. henryi*, *C. mollissima*, *C. pumila*, *C. sativa*, and *C. seguinii* (2) – are distributed mostly in the temperate regions of the world and with species native ranges in north America, Europe and Asia. Historically, these trees played critical roles in facilitating expansion and settlement of human populations into new territories and thus, in many regions of the world, these trees are designated heritage trees under special laws of protection to insure their survival.

Unfortunately, as with many of our forest trees, the continued survival of these dominant forest tree species is continually challenged by problems of forestland reduction, overharvesting, invasive pests/ pathogens, and rapidly changing environmental conditions. Addressing these challenges proactively with knowledge-based strategies to sustain and improve the health of our forest tree resources increasingly relies on having highly characterized diverse germplasm materials, highly developed genetic and genomic tool resources in key species and state of the art breeding programs for capturing and mobilizing important traits (e.g. pathogen resistance) into improved tree materials for replant and restoration efforts.

*Castanea* species are ideally suited for addressing fundamental questions on the nature of host/pathogen genome coevolution and invasive pathogen biology in trees. For example, Chinese chestnut (*C. mollissima*) has coevolved with and has resistance to two major invasive pathogens*, Cryphonectria parasitica* and *Phytophthora cinnamomi*, both responsible for the complete demise of the susceptible American chestnut (*C. dentata*) as a dominant US forest species after their unintentional introduction to the US from Asia (3,4). Chinese chestnut and American chestnut can hybridize, and the resulting interspecies hybrid families segregate for resistance as well as other traits of interest (morphological and phenological traits). Additionally, it is possible to obtain early flowering in these trees (within a year) substantially reducing generation time for basic and applied genetics approaches in heritage forest tree restoration programs (5). In this context, these species afford an excellent opportunity to advance our genetic understanding of the coevolution of host/pathogen complexes in forest trees and other traits as well. We have focused on the development of genetic and genomic resources in Chinese chestnut as a key tree species in *Castanea* that is currently used in several breeding programs as a donor of resistance to *C. parasitica* and *P. cinnamomi* in American chestnut and other important traits. (1,6). Here, we present the development of a Chinese chestnut whole genome sequence and its implementation in studies to understand the diversity and evolution of important host/pathogen complexes and other traits important in adaptation and response to climate change required for restoration of threatened chestnut species.

## RESULTS

### The Chinese chestnut genome assembly and structural features

Our goal to develop a high-quality, chromosome-scale genome for the *C. mollissima* cultivar Vanuxem, proceeded through several rounds of *de novo* assemblies and sequence anchoring to a reference genetic map. An initial, *de novo* assembly of the genome, version V1.1, was produced in 2013 and released to the public in January 2014 as a browser and searchable database at the Hardwood Genomics website (www.hardwoodgenomics.org). A total of 13.7 Gb of 454 technology data (26.2 million reads) plus 46 Gb of Illumina MiSeq data (from 149.6 million reads) was produced. An optimal, hybrid *de novo* assembly was selected using the heterozygosity option in Newbler v2.8 assembly software. The assembly placed 724 Mb in 41,260 scaffolds, which provided 91.2% overall coverage of the Chinese chestnut genome (estimated at 794 Mb by flow cytometry) (7), with an N50 scaffold length of 39,580 bp, an L50 of 5,019 scaffolds, and largest scaffold at 429,344 bp. The V1.1 scaffolds included 27,264 gaps, with an overall gap length of 13.5 Mb. A total of 36,478 gene models, and 38,146 transcripts and peptide sequences were predicted and annotated in the V1.1 assembly, which were also included for public access at the Hardwood Genomics database.

To better support basic research and genome-wide-selection models for disease-resistance breeding, we then focused efforts on building a more contiguous assembly and anchoring sequences to chromosomal locations. Initially, contigs were merged based on co-localization with BAC-end sequences in the *C. mollissima* physical map (8) and gaps closed manually. The resulting Version 2 (V2) assembly increased to 760 Mb, consisting of 60,546 contigs and 14,358 scaffolds (N50 2.75Mb). Further rounds of contig merging and gap closing using PACBio reads produced an improved hybrid assembly (V3.2) of 14,110 contig sequences spanning 725.2 Mb, with maximum and minimum contig sizes of 663Bb and 2 kb, respectively, with no internal gaps and minimal sequence ambiguities. Mapping of DNA marker sequences from the integrated *C. mollissima* physical and genetic map (9) indicted that the V3.2 assembly accomplished close to complete coverage of the genome (98-99%). For detailed descriptions of assembly versions, see Supplementary Table S1).

The improved *de novo* assembly served as the starting point for assembling pseudomolecule sequences to represent each of the 12 chestnut chromosomes (hereafter referred to as “pseudochromosomes”). Initially, contigs were anchored to a higher marker-density version of the chestnut research community’s reference genetic linkage map for the Vanuxem genotype (9). To increase the number of anchored contigs, we also used DNA markers from three additional *Castanea* genetic maps, in regions where those maps were consistent with the initial framework assembly. At that point, the anchoring results provided evidence of high collinearity between oak and chestnut chromosomes (see “Analysis of genome structure” section below). Thus, we also used a dense *Quercus robur* genetic map (Bodenes, INRA Pierroton, unpublished results) to anchor contigs in places of colinearity where a chestnut marker had not been found. This approach resulted in 11,795 markers at 4,618 unique genome positions used to anchor and order 4,403 unique sequence contigs. After limiting contigs anchored to multiple chromosomal locations to the single most robust position, a final total of 4,099 of the contigs were positioned relative to their positions on the community’s reference genetic linkage map (9), yielding 12 pseudochromosome sequences ranging from 25.9 Mb (LG_L) to 60.2 Mb (LG_A), and totalling 421.3 Mb. This represents only 53% of the estimated total genome length of 794 Mb.. Gene content analyses of the pseudochromosomes revealed that (67%) of 30,835 putative gene models in Chinese chestnut (see functional features section) were present in these pseudochromosomes. The chromosome-naming strategy was based on maintaining the long-standing linkage group lettering convention within the chestnut research community. A summary of the V4.0 pseudochromosome assembly statistics is provided in Table 1.

**Table 1.** Chinese Chestnut pseudo-chromosome structural features.

| METRIC | Assembly V 4.0 (chromosome-anchored assembly) |
| --- | --- |
| **Contig Assembly** | • **421,3Mb** (4,099 contigs) from *de novo* assembly V3.2 anchored to *C. mollissima* chromosomes<br>• Anchored sequence per chromosome ranges from 25.9Mb (LGL) to 60.2 Mb (LGA)<br>• 303.9Mb (10,011 contigs) unanchored contig sequences<br>• 57.8Mb estimated gaps (based on estimated full genome length of 794Mb) |
| **Gene models:** | • 30,832 high quality gene models<br>• 20,770 high quality gene models contained within the contigs anchored to chromosomes |
| **Validations:** | • BUSCO reported 1,355 of 1,440 expected single-copy genes are complete and present within the *C. mollissima* genome<br>• Alignments of BAC end sequences from the *C. mollissima* physical map confirmed genetic map-based order of contigs in the scaffolded chromosomes |

Validation of the completeness of assembly of the gene space was obtained by BUSCO analysis, which accounted for 93% of expected single copy genes in the V3.2 assembly. The overall placement of anchored contigs was supported by query of the pseudochromosome sequences with BAC-end sequences from the Vanuxem physical map tiling path. A summary of the V4.0 pseudochromosome assembly statistics is provided in Table 1.

A website for the chestnut genome versions 1.1, 3.2, and 4.0, as well as the chestnut blight QTL sequences, was constructed at https://hardwoodgenomics.org/content/genomic-data. The website contains links to download the whole genome contigs, scaffolds, and pseudo-chromosomes as well as predicted genes, transcripts, Open Reading Frames, and proteins and annotations, as well the QTL contigs and scaffolds and their gene content. J-Browse implementations for the whole genome and QTLs and associated analyses are located at the URLs https://hardwoodgenomics.org/tools/jbrowse/?data=chinese_chestnut and https://hardwoodgenomics.org/tools/jbrowse/?data=chinese_chestnut_qtls. The sequences are also available at the NCBI BioProject No.

### Repetitive landscape in chestnut

A total of 1,925 sequences were identified as repetitive elements within the *C. mollissima* assembly. Excluding rRNAs, the repeat sequences totaled 369,149,927 base pairs, or just over 50% of the genome, using the combined RepeatModeler and RepBase Plant Library and with a query species of “eudicotyledons”. As shown in detail in Supplementary Table S2, most repetitive elements in chestnut are interspersed repeats. The largest class of the repetitive elements were “unclassified” (25.09%), with the second most abundant class being retroelements (21.05%). Within retroelements, the most abundant form of repetitive element was long terminal repeats (LTRs), consisting of 17.48% of the total genome. The most prevalent DNA transposon family identified was “Hobo-Activator.”

### *In situ* assignment of pseudochromosomes to chestnut chromosomes

Individual LGs were assigned to specific chestnut chromosomes by fluorescent *in situ* hybridization (FISH). From 2 to 8 markers per linkage group were chosen from the set of mapped markers on the Chinese chestnut reference genetic map (9) that had been used to integrate the linkage groups from top to bottom on the Chinese chestnut BAC physical map (8). In addition, ribosomal DNA (18S–25S and 5S rDNA) probes were used to identify their LG-specific cytological positions. For each linkage group and the corresponding marked regions of the physical map, BACs were chosen as probes for FISH on chestnut root tip chromosome spreads. (see Supplementary Table S3 for a full list of BAC clones selected). Since primary constrictions serve as cytologically visible landmarks for centromere position, we were able to anchor the linkage groups to their respective chromosomes and determine the relationship of the linkage group to the long and short arms of each corresponding chromosome (Fig. 1). The zero cm linkage map position was found to be associated with the short arm of nine chromosomes and the long arm of three chromosomes (LGs C, G and L). The cytological analyses enabled a putative designation of six of the twelve Chinese chestnut LG-specific chromosomes (LGs A, B, C, F, G and I) as metacentric and/or near metacentric, four (LGs E, H, J and K) as near sub-metacentric and two (LG D and LG L) as clearly sub-metacentric chromosomes. Of the 54 BAC clones and two ribosomal DNA probes (18S–25S rDNA and 5S rDNA) used in FISH, the cytological positions (i.e., orientations) of all but three BAC clones were concordant with their expected linkage group position on the genetic map (Fig. 1). We observed the major 18S–25S rDNA distally on the short arm of LG_H chromosome, but not the previously reported minor second locus (10). A satellite (SAT) knob and nucleolus organizer region (NOR) were observed on the LG_H chromosome where the BAC H-C5 clone hybridized proximally to the 18S–25S site (Fig. 1). One 5S rDNA site was located in the middle of the short arm of the LG-E chromosome. Representative cytological images showing examples of multiple BAC probe assignments by FISH to the *C. mollissima* LG_D chromosome is shown in Fig. 2, along with the corresponding locations of markers on the reference genetic linkage map.

**Figure 1.**
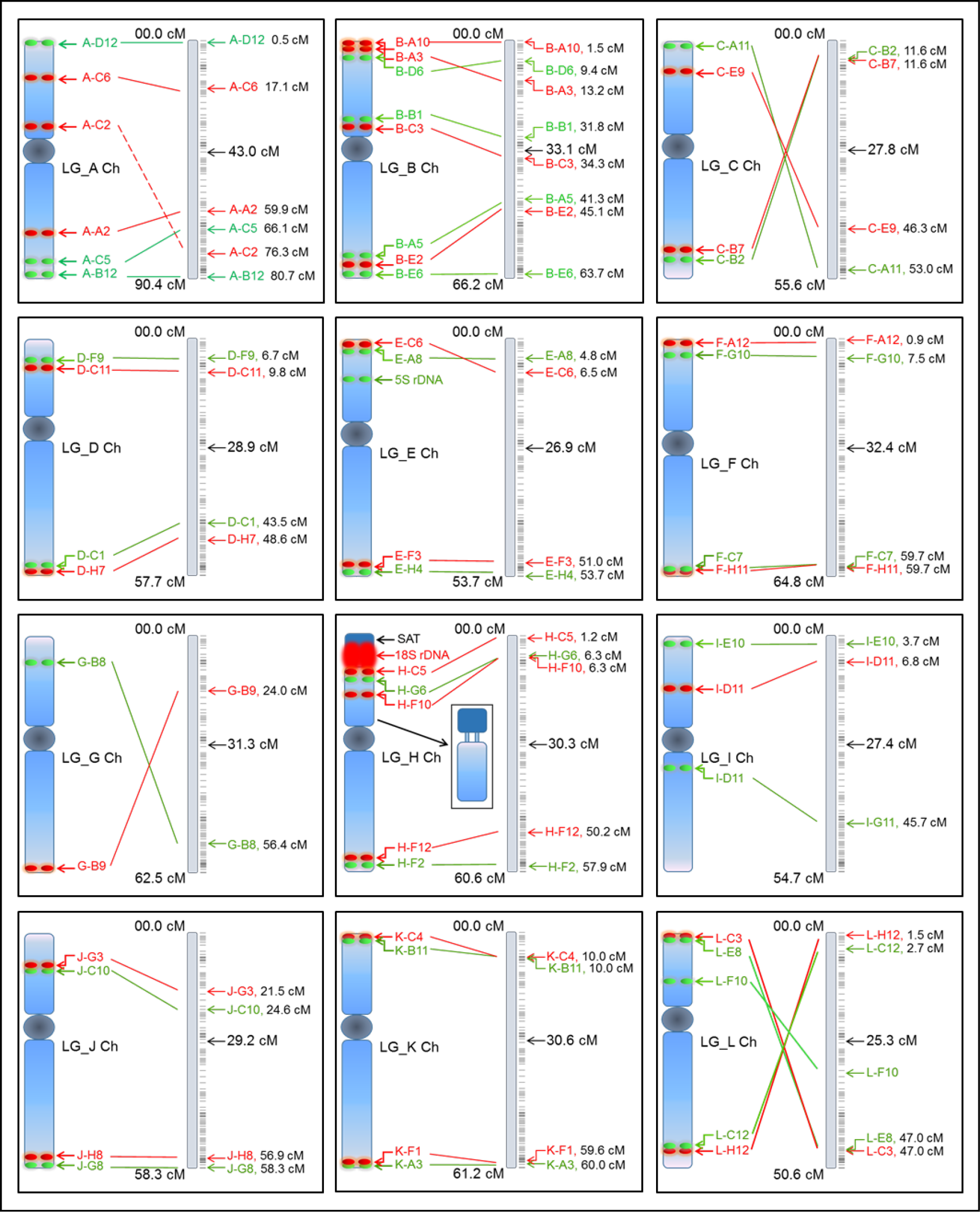
Diagrammatic representation of BAC-FISH mapping results for assignment of *C. mollissma* chromosomes to their corresponding linkage groups. The putative position of the centromeres of each LG map is delineated. The insert in the panel of LG_H shows the position of the satellite and rDNA expansion in the LG_H chromosome.

**Figure 2.**
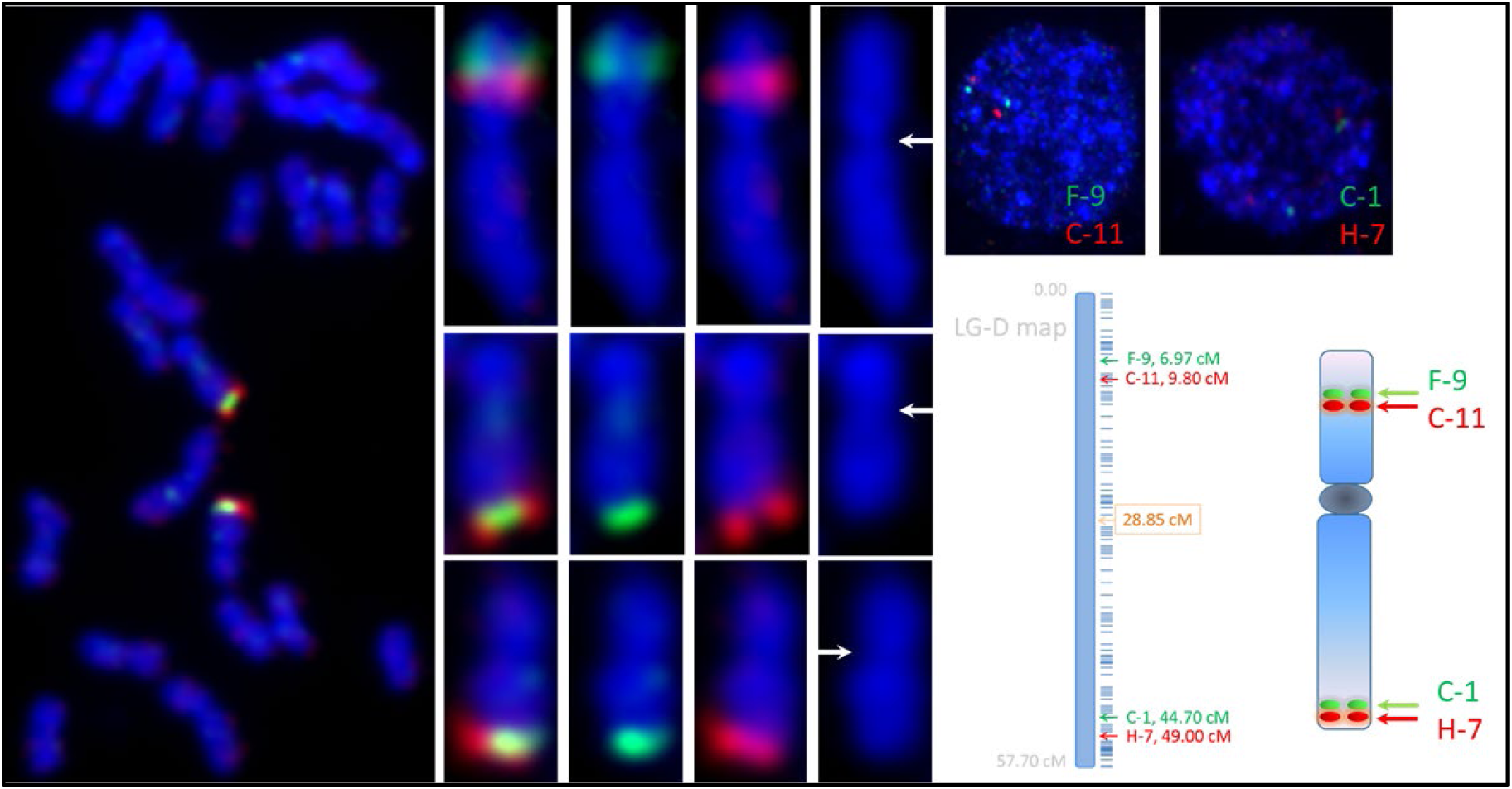
Example of multiple-probe BAC-FISH mapping result for assignment of *C. mollissima* chromosomes Linkage group D to its corresponding linkage group. FISH was conducted with four selected BAC clones (two from opposite ends of each arm) on Chinese chestnut chromosome spreads: a) a complete root tip metaphase chromosome spread showed two BAC FISH signals (BAC C1, 44.70 cM, green signal; BAC H7, 49.00 cM, red signal) located on the long arm of the LG_D homologs (chromosomes); b1-4) an enlarged FISH image of a pro-metaphase chromosome with BAC clones (BAC F9, 6.97 cM, green signal; BAC C11, 9.80 cM, red signal) located on the short arm of the same chromosome; b1, a superimposed image from DAPI (blue chromosome background), FITC (green signal), and Cy3 (spectrum-orange/red signal) filters; b2, an image from DAPI and FITC filters; b3, an image from DAPI and Cy3 filters; b4, an image from DAPI filter, and it is the same for c1-4 and d1-4; and c1 and d1, enlarged images of the homologous pair of the LG_D metaphase chromosomes. The white arrows in b4, c4, and d4 showed the primary constriction (i.e., the centromere); e) and f) are from two interphase FISH nuclei showing the BAC FISH signals; g) a diagrammatic representation of the LG_D map; and h) a diagrammatic representation the LG_D chromosome delineated by the primary constriction (centromere) and showed the short (S) arm and long (L) arm with respective BAC FISH signals.

### Chinese chestnut genome functional features

#### Annotation Statistics and Quality Assessment

BRAKER2, which is a combination of previously established gene prediction tools Genemark-EX and AUGUSTUS, was used for gene predictions in conjunction with RNA-Seq and protein homology information. The statistics of alignments to the V3.2 genome assembly of RNA-Seq reads from several transcriptome projects can be found in Supplementary Table S4. Overall alignment of RNA reads to the assembly was high. From the alignment results, GenomeThreader predicted that 24,559 genes would be present in the Chestnut assembly. The BRAKER gene-finding algorithm (11) predicted a total of 50,911 genes for the assembly before filtering steps were taken. A manual, evidence-based filtering protocol for genes supported by RNA-Seq and GenomeThreader gene models yielded 30,835 genes, which is representative of gene number estimates for other Fagaceae species, *Q. suber*, 37,724 genes (12); *Q. robur*, 25,808 genes (13).

Of these remaining genes, 2,085 were supported by GenomeThreader only, 16,231 were supported by RNA-Seq only, and 12,518 were supported by both RNA-Seq and GenomeThreader. All of the predicted gene model names associated with each contig and with each Linkage Group are provided at www.hardwoodgenomics.org.

As a check on completeness of the annotation, the predicted genes were compared to single-copy orthologs found in the group Embryophyta, using the BUSCO python script (14). BUSCO analysis reported that 1,355 of the 1,440 expected single-copy genes were complete and present within the *C. mollissima* genome. Of the complete BUSCOs, 1,266 were single-copy, with the remaining 89 present in more than one location in the genome. Of the unaccounted single-copy orthologs, 33 were fragmented ORFs and 52 were missing. This result of all but a few expected of embryophyte single-copy genes being present in the genome and complete, indicates that our assembly and annotation and annotation of the gene space is largely complete.

#### Functional Annotations

As a first step in evaluating the completeness of the gene resources in our assembly, we compared the shared orthogroups for the 30,835 chestnut gene models among a selection of model species with complete genome resources representing both woody, tree and herbaceous plants (Table 2). Orthologous groups of chestnut proteins were identified using OrthoFinder2 and clustered with known Arabidopsis, peach, poplar, and grape orthogroups. OrthoFinder2 placed 71.4% of 163,425 predicted proteins from the 5 species into 16,687 orthogroups (for detailed results see Supplementary Table S5), with a mean size of 7 proteins. Only 212 species-specific orthogroups were obtained, while 11,624 orthogroups had representatives in all 5 species.

**Table 2.** Orthogroups shared among species (numbers of protein overlaps)

| Species | Arabidopsis | Chestnut | Peach | Populus | Grape |
| --- | --- | --- | --- | --- | --- |
| Arabidopsis | 13,178 | 12,425 | 12,670 | 12,738 | 12,484 |
| Chestnut | 12,425 | 15,044 | 13,976 | 14,022 | 13,901 |
| Peach | 12,670 | 13,976 | 15,084 | 14,224 | 13,961 |
| Populus | 12,738 | 14,022 | 14,224 | 15,297 | 14,083 |
| Grape | 12,484 | 13,901 | 13,961 | 14,083 | 15,027 |

The analysis showed that the chestnut genome reference, as judged by number of shared orthogroups, does not significantly differ from other closely and more distantly related species. Much of the historic interest in the genetics in chestnut has focused on resistance to the invasive pathogens that eliminated American chestnut as a dominant species of the eastern forests of North America. For this reason, we were particularly interested in the potential discovery of genes that underlie resistance to fungal or oomycete pathogens. The phenylpropanoid pathway has been shown in numerous studies [e.g. avocado (15), eucalyptus (16)] to underpin stress response in trees to biotic and abiotic stressors to chestnut and thus was of direct interest to this study (see Table S5 and Table 3). Additional results of functional annotations from InterProScan for the chestnut gene models are provided at www.hardwoodgenomics.org.

**Table 3.** Predicted numbers of lignin monomer pathway orthologous gene models in Chinese chestnut and model tree genomes.

| gene family | Orthogroups | Arabidopsis | chestnut | peach | Populus | grape |
| --- | --- | --- | --- | --- | --- | --- |
| PAL | 1 | 3 | 3 | 2 | 4 | 12 |
| C4H | 2 | 1 | 2 | 2 | 3 | 3 |
| 4CL | 10 | 11 | 14 | 13 | 17 | 14 |
| HCT | 1 | 1 | 1 | 2 | 2 | 1 |
| C3H | 1 | 1 | 2 | 4 | 3 | 1 |
| CCoAOMT | 4 | 6 | 11 | 7 | 5 | 9 |
| CCR | 6 | 8 | 8 | 9 | 11 | 11 |
| F5H | 1 | 2 | 1 | 2 | 3 | 3 |
| COMT | 10 | 10 | 16 | 28 | 21 | 28 |
| CAD | 5 | 7 | 17 | 18 | 11 | 17 |

#### The NBS LRR Gene family

A Pfam search using NBS and LRR motifs on the orthogroups above from the peach, grapevine, poplar, and oak genomes and the chestnut pseudochromosome high quality, supported gene model produced the following NBS-LRR gene family totals:

*Chestnut: 300; Peach: 386; Vitis: 450; Poplar: 556; Oak: 874*.

Our parameters for the pfam search produced different totals than previously reported for peach, grapevine, poplar and oak. However, this result is consistent with the general observation that this gene family has experienced a major expansion in *Q. robur* (13). In contrast, the NBS-LLR family of disease-resistance genes in the Chinese chestnut genome appears to be reduced to a number even lower than in the comparatively small peach genome [265 Mb, (17]. There are a number of possible reasons including– 1) tandemly repeated genes such as NBS-LRR were not anchored well using the genetic linkage map markers, which only represent unique sequences in the genome, 2) high heterozygosity levels in the chestnut genome coupled with high sequence similarities among NBS-LLR genes may have caused the heterozygosity-option in the Newbler software to collapse the copy number of tandemly repeated genes during the assembly process.

#### Analysis of genome structure

Since whole genome sequences are available for a several tree species, it was of interest to assess the level of genome preservation between chestnut and other species that have significant information on gene/trait associations. If the preservation of genome organization is high enough, this gene/trait information can potentially be translated across species and thus leverage the resources invested in one species to assist in knowledge gain in another. In this regard, we performed genome comparisons by alignments of chromosomes between chestnut and oak (*Q. robur*) which along with chestnut is within the Fagaceae family (Fig. 3), between chestnut and peach (*P. persica* Batsch.) (Fig. 4), a member of the Rosaceae for which there is rapidly increasing information on gene/trait associations (18). Overall, the Circos plots in figures 3 and 4 illustrate the high degree of macro-synteny at the whole chromosome level between chestnut and oak, in genome structure among species in the Fagaceae family that have been previously reported from genetic mapping studies (7, 9, 12, 13). Figure 3 also reveals some divergence in gene order or gene copy number at a finer-scale between chestnut and oak. Major blocks of synteny were also observed between the chestnut and peach genomes (Fig. 4). This illustrates, as previously reported for oak (13), that only a few chromosomal breaks and fusions may account for the differences in overall genome organization between the Fagaceae and Rosaceae families from their last common ancestor. A more detailed illustration of macro- and micro-syntenies for individual chestnut chromosomes with the oak and peach genomes are shown in Supplemental Figures S1 and S2. The individual alignments of chestnut chromosomes revealed that the chestnut pseudochromosomes do contain substantial numbers of dispersed repetitive DNA elements shared with, and also widely distributed across, the oak genome (Fig. S1). Fewer such widely dispersed elements appear to be shared between the chestnut chromosomes and the peach genome (Fig. S3). However, the individual chestnut chromosomes alignments reveal the rearrangements in genome organization at the micro-level at much greater resolution.

**Figure 3.**
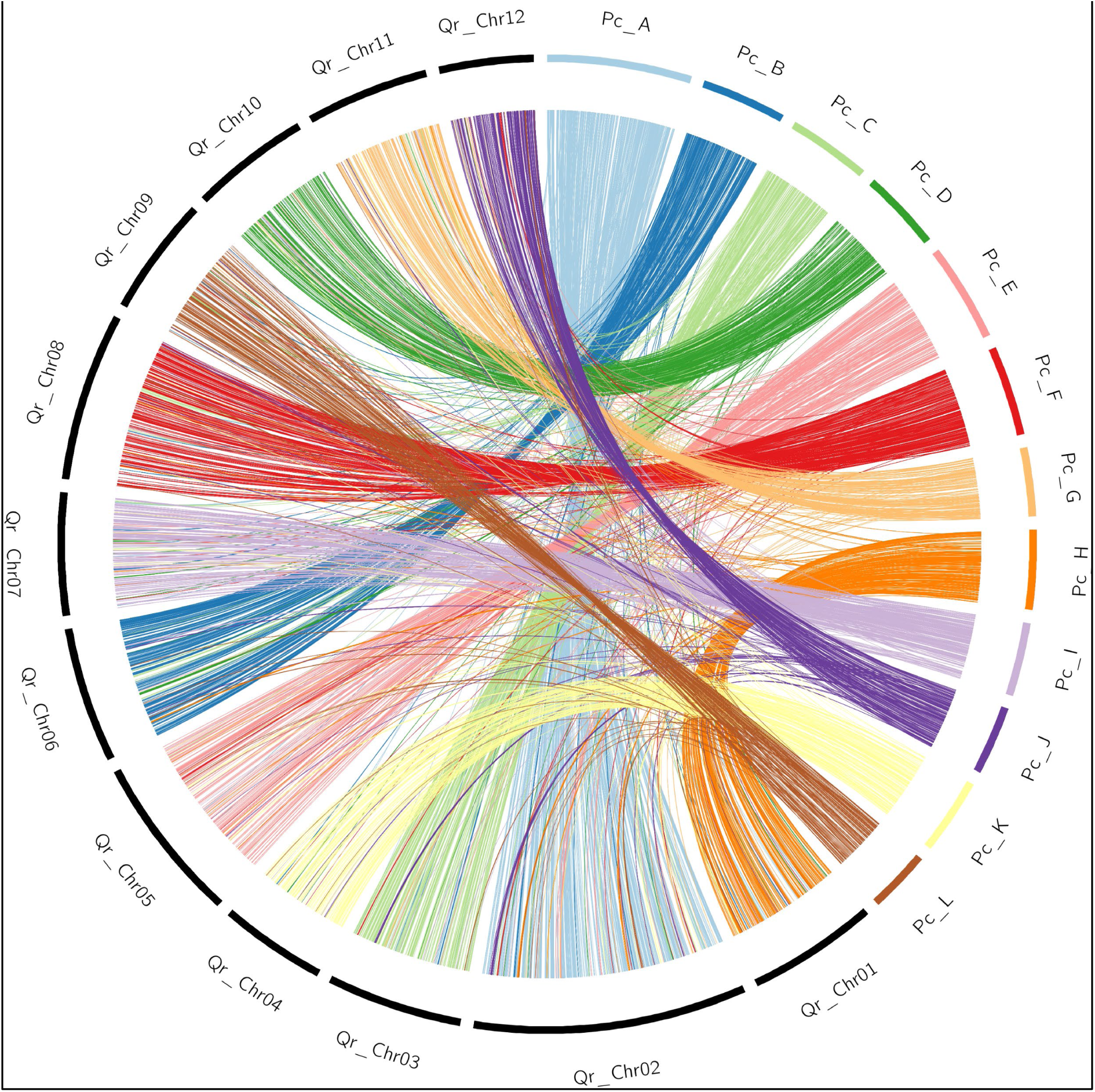
Circos plot of alignments of orthologous genes in C. mollissima pseudochromosomes (Pc_A-L) vs Q. robur chromosomes (Qr_1-12). Alignment of the genomes followed filtering to reduce the number of ortholog repeats. Pc, pseudochromosome. *C. mollissima* pseudochromosome naming convention of adheres to genetic linkage group (13) assignments.

**Figure 4.**
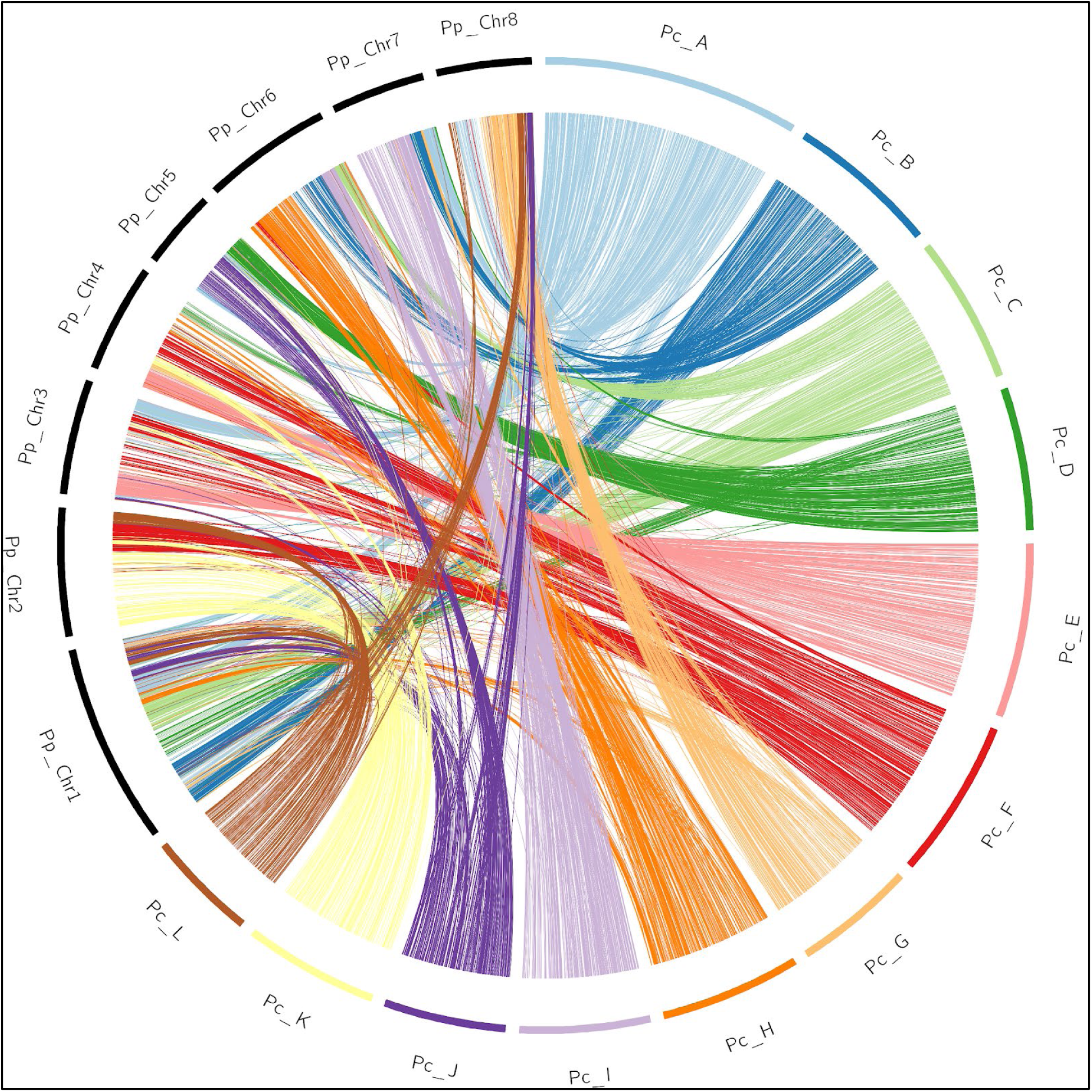
Circos plot of alignments of orthologous genes in C. mollissima pseudochromosomes (Pc A-L) vs P. persica chromosomes (Pp_1-8). Alignment of the genomes followed filtering to reduce the number of ortholog repeats. Pc, pseudochromosome. *C. mollissima* pseudochromosome naming convention of adheres to genetic linkage group (13) assignments.

### Castanea comparative genomic analyses

The availability of a reference *C. mollissima* whole genome sequence has enabled the analyses of species of diversity within the genus and the potential impacts of host/pathogen co-evolution in *Castanea* species genomes, both genome wide and the level of individual mapped resistance-conferring QTL intervals.

To assess genetic variation and heterozygosity among Castanea species endemic to China, we sequenced 43 trees of *C. mollissima* across 34 localities, 28 trees of *C. seguinii* across 23 localities and 27 trees of *C. henryi* across 19 localities spanning their geographic ranges (Fig 5), using the Illumina Genome Analyzer (HiSeq 2500) with a read length of 150 bps. This generated a total of 13.4 billion short reads and an average of 136.4 million short reads (25.57× genome coverage) per tree. After filtrating low-quality reads, alignment and genotyping individually, we identified a total of 66.03 million single nucleotide polymorphic (SNP) sites in 98 trees of three species. The average heterozygosity was higher in *C. seguinii* (0.0055 ± 0.00098) and *C. henryi* (0.0052 ± 0.00067), than that in *C. mollissima* (0.0039 ± 0.00029). In the dataset without missing bases, a total of 13.73 million SNPs were retained and the number of SNPs *in C. seguinii, C. henryi* and *C. mollissima* were 7.36, 5.92 and 3.90 million, respectively; and the number of SNPs with minor allele frequency (MAF) > 5% were 3.90, 2.58 and 1.83 million.

**Figure 5.**
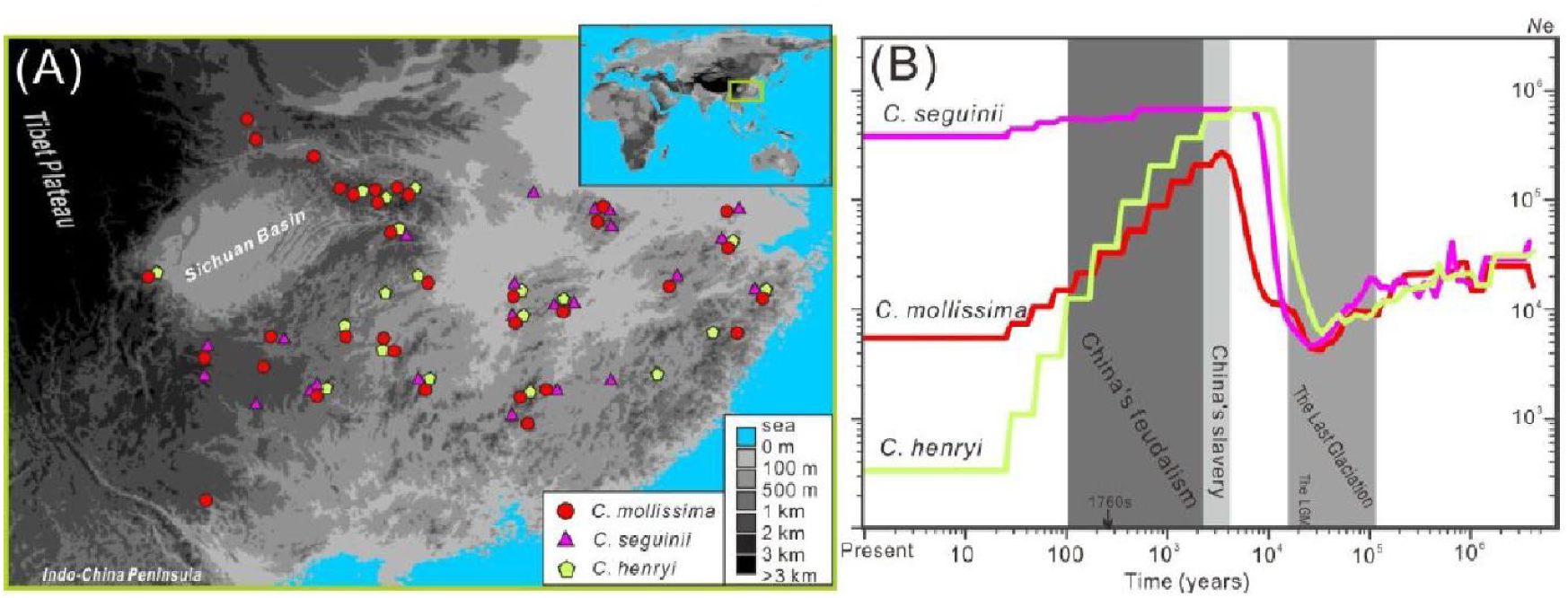
Source populations. A) Locations of all samples; B) Profile of changes in effective population sizes for the three species. Color scheme: red, C. mollissima; fuchsia, C. seguinii; green, C. henryi.

To validate the quality of our SNP calls, 4 trees of *C. mollissima* and 3 trees of *C. henryi* were re-sequenced in parallel. These short reads for validation were processed in identical pipeline with the processing from quality control to genotyping blind to the fact that they were duplicates. Then we compared the genotypes in the two duplicated samples for each of 7 trees. The rate of discordant genotypes between the two duplicate samples was very low (0.66 x10^−3^, 0.63 x10^−3^ 0.67 x10^−3^, 0.94 x10^−3^, 0.70 x10^−3^, 0.74 x10^−3^, 0.75 ×10^−3^ per SNP) for each tree, reflecting an average error rate lower than 0.37 ×10^−3^ per SNP (∼ 34.32 in Phred-logarithmic scale). It suggested that the SNPs produced by the present sequencing strategy and filtration protocol are of high quality.

If blight infection (*C. parasitica*) is a major factor in shifting population sizes of *Castanea* species, as shown in American chestnut forest (19), the three *Castanea* species native in China would show synchronous recent reductions of effective population size (*N*e). We inferred the historical effective population sizes (Ne-s) of *C. seguinii, C. mollissima* and *C. henryi*, using the model of sequential Markov coalescent with population samples, SMC++ (20). These analyses showed well-defined demographic histories for the three species from one million to tens of years ago (Fig 6).

**Figure 6.**
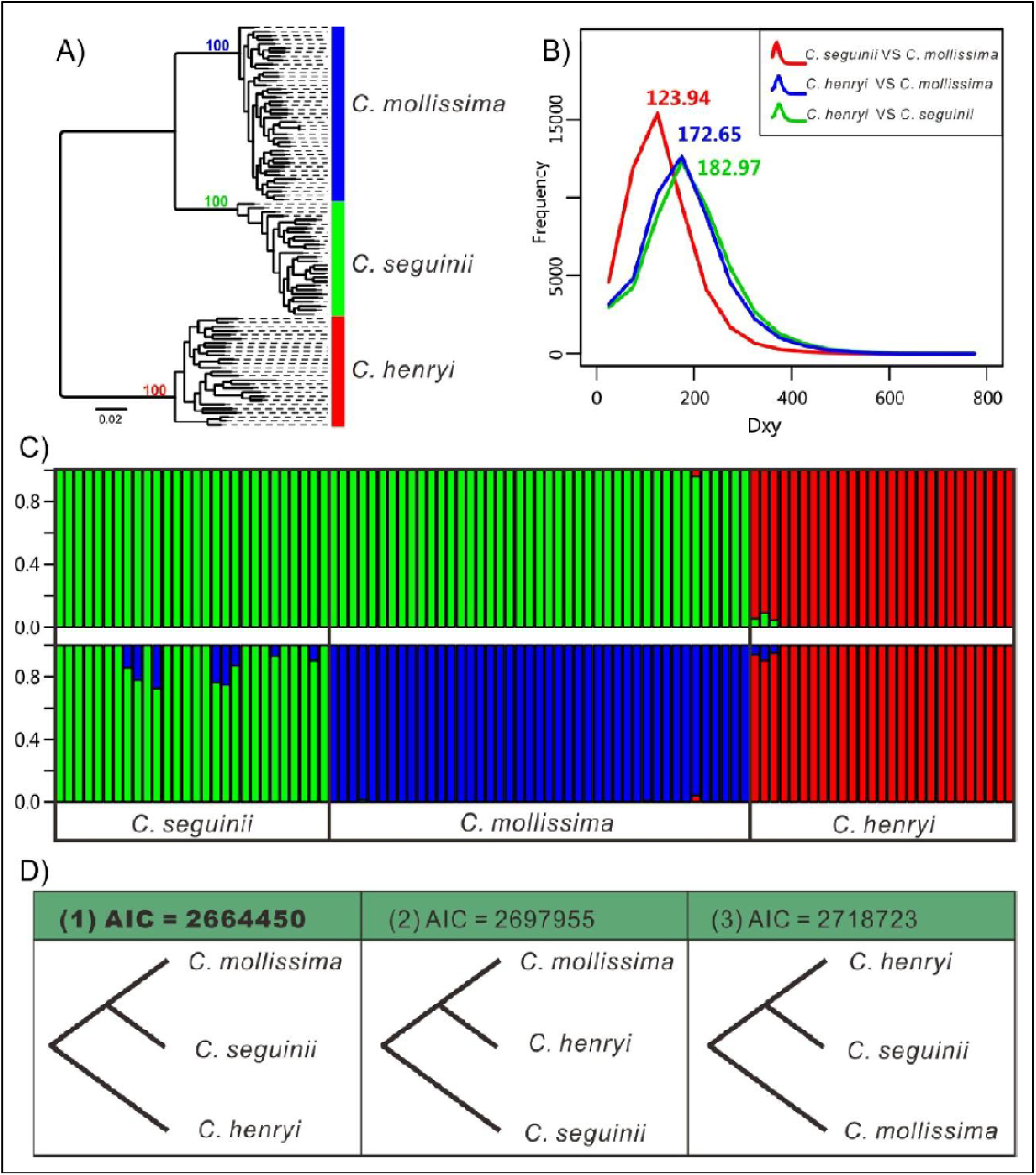
Well-defined demographic histories for *C. seguinii, C. mollissima* and *C. henryi*. A) Genealogical tree of all 98 samples, B) the absolute genetic differentiation between each pair of species, C) ancestry assignments, D) coalescent test of species tree.

The high blight resistance within *C. mollissima* and *C. seguinii* may be due to tolerance in their common ancestor and/or introgression of resistance genes. To distinguish these two likely causes, we performed: 1) a sliding-window (20 Kb size) individual clustering analyses of absolute divergence along the genome employing a phylogenetic tree based on the concatenated sequences; 2) an admixture analysis; and 3) a coalescent test. All of them supported that *C. mollissima* and *C. seguinii* constituted a single clade (Fig 6).

The close relationship between *C. mollissima* and *C. seguinii* and their distribution pattern may suggest past hybridization, which might play a role in the evolution of blight resistance within the Chinese chestnut. The transfer of small genomic regions from a donor species into a recipient species, is characterized by smaller divergence time at the introgressed locus, relative to the time of divergence from the donor species. To test if introgression between *C. mollissima* and *C. seguinii* had occurred, we introduced a modified dXY statistic, called *tit*, which reduced the false positives caused by heterogeneity of the mutation rate along the genome resulting from the use of the genomic sequences of *C. henryi*. We identified 172 genes with significantly lower *tit* than expected from coalescent simulations, after correcting for the effects of possible strong selection in their ancestral population. Low levels of *tit* can also be explained by the effects of conservative evolution at some genes. We used the nucleotide diversity π to infer the lineage-specific conserved genes which exhibit a low level of polymorphism along the genomes of *C. mollissima* and *C. seguinii* and 60 conserved genes were identified with a false positive rate 5%. Thus, our analysis suggested that at least 112 genes were potentially involved in interspecific gene exchange between *C. mollissima* and *C. seguinii*. The HKA test further supported that 107 putatively introgressed genes showed significant signatures of selection in either *C. mollissima* or *C. seguinii*, indicative of likely a possible adaptive introgression between them.

#### Signatures of selection in the blight resistance QTL *cbr1* region on Linkage Group B in *C. mollissima* vs *C. dentata*

The relative ease of hybridizing *C. mollissima* and *C. dentata* enabled a number of QTL mapping studies (9, 21), that provided information on the genetics of blight disease resistance in chestnut. With the availability of the *C. mollissima* genome, we were interested to know if genes in the mapped QTL regions had differing signatures of selection between American chestnut and the Chinese chestnut, supporting the hypothesis that the blight pathogen had exerted selection pressure on the Chinese alleles for these genes over the period of its coevolution of the host.. For the identification of selective sweep regions in the Chinese and American chestnut genomes, we focused on Linkage group B where multiple mapping studies had demonstrated a significant QTL for blight resistance. Statistical tests for neutrality were calculated for two re-sequenced pools (five *C. dentata* and five *C. mollissima* genotypes) of American and Chinese chestnut. Two statistical parameters were calculated for each species: pooled nucleotide diversity (*Pi*) and pooled Tajima’s D (TajD). A 5 kb sliding window was employed to allow gene level resolution. The distribution of transformed nucleotide diversity in American (*Pi Cden*/*Pi Cmol*) and Chinese (*PiCmiol/PiCden)* chestnut pools plotted along the *C. mollissima* v3.2 pseudochromosome 2 (LG_B) are shown in Fig. 7A. Nucleotide diversity in *C. mollissima* pool aligned against the *C. mollissima* reference genome was lower than that in the *C. dentata* pool. In total 8 genomic regions (5kb and larger) were detected as candidate regions (CKR) under purifying selection in the *C. mollissima* pool.

**Figure 7.**
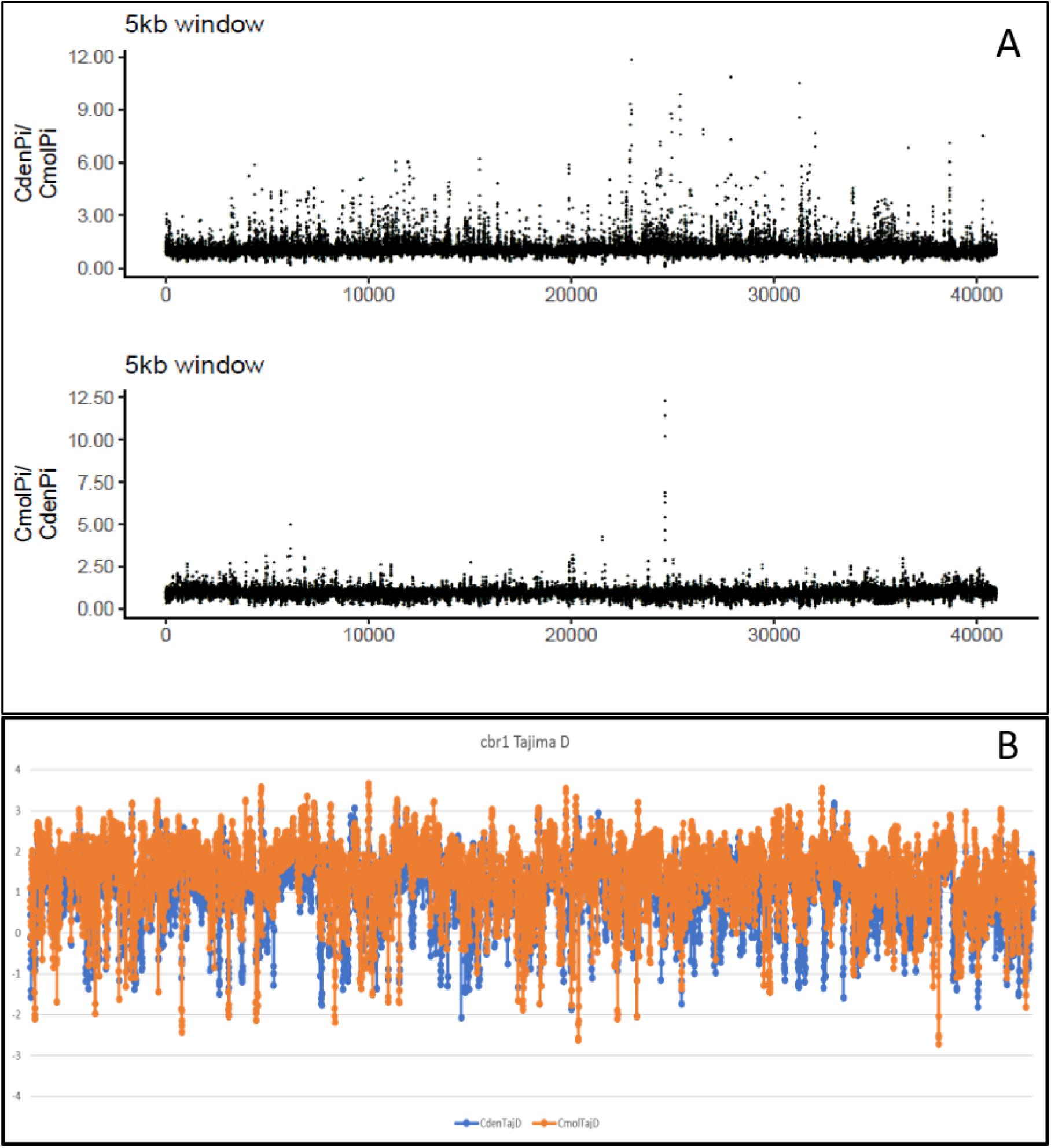
Comparisons of nucleotide diversity and TajD values across the entire cbr1 *Cryphonectria* resistance QTL region on LGB. **A.** Distribution of transformed nucleotide diversity ratios in American (*Pi Cden*/*Pi Cmol*) and Chinese (*PiCmiol/PiCden)* chestnut pools plotted along C. mollissima (Vanuxem) for pseudochromosome B (LG_B). A dashed horizontal line indicates the cut-off used for identification outliers representing top 0.5%. **B.** A profile of TajD values determined from nucleotide diversity ratios across the cbr1 *Cryphonectria* resistance QTL region on LGB as bracketed by DNA markers in the reference *C. mollissima* genetic linkage map (9). Each point represents results for a specific sliding window across the QTL. Orange dots and lines are values for *C. mollissima*; Blue dots and lines are values for *C. dentata*. Values below 0 show sequences that may be under greater positive selection for that species. Values lower than −1.5 are considered strong signatures of selection but must be compared to the value in that window for the other species to know if that selection is differential or not, versus the other species, which is difficult to assess at this condensed horizontal axis scale.

We focused on Linkage Group B for this assessment, as the blight-resistance QYL cbr1 on LGB is the most consistent among mapping studies. The QTL located on LGB with the enhanced genetic linkage map family (22) overlapped with but was not entirely syntenic with cbr1 QTL determined with the original, smaller reference linkage mapping family (9). Assuming that this region of overlap is particularly important in blight resistance, we then focused on TajD profiles in the LGB QTL region of overlap from the two mapping studies (9, 22). In this overlap region, we detected only one highly significant signature of selection (TajD <-2) in this region, in sequence contig 0002011 (Fig. 8). The 002011 contig contains 14 open reading frames. The strong Tajima’s D value for signature of selection overlaps with the predicted ORF #13, which has very strong BLASTn and BLASTp alignment scores (Evalues of 0 and 75-96% identities) to Type I Inositol Polyphosphate 5-Phosphatase 1 genes in woody plants. UniProt describes the IP5P1 protein as “involved in the abscisic acid (ABA) signaling pathway (PubMed:12805629).” The IP5P1 gene’s top GO functional category is Biological process for “abscisic acid-activated signaling pathway”.

**Figure 8.**
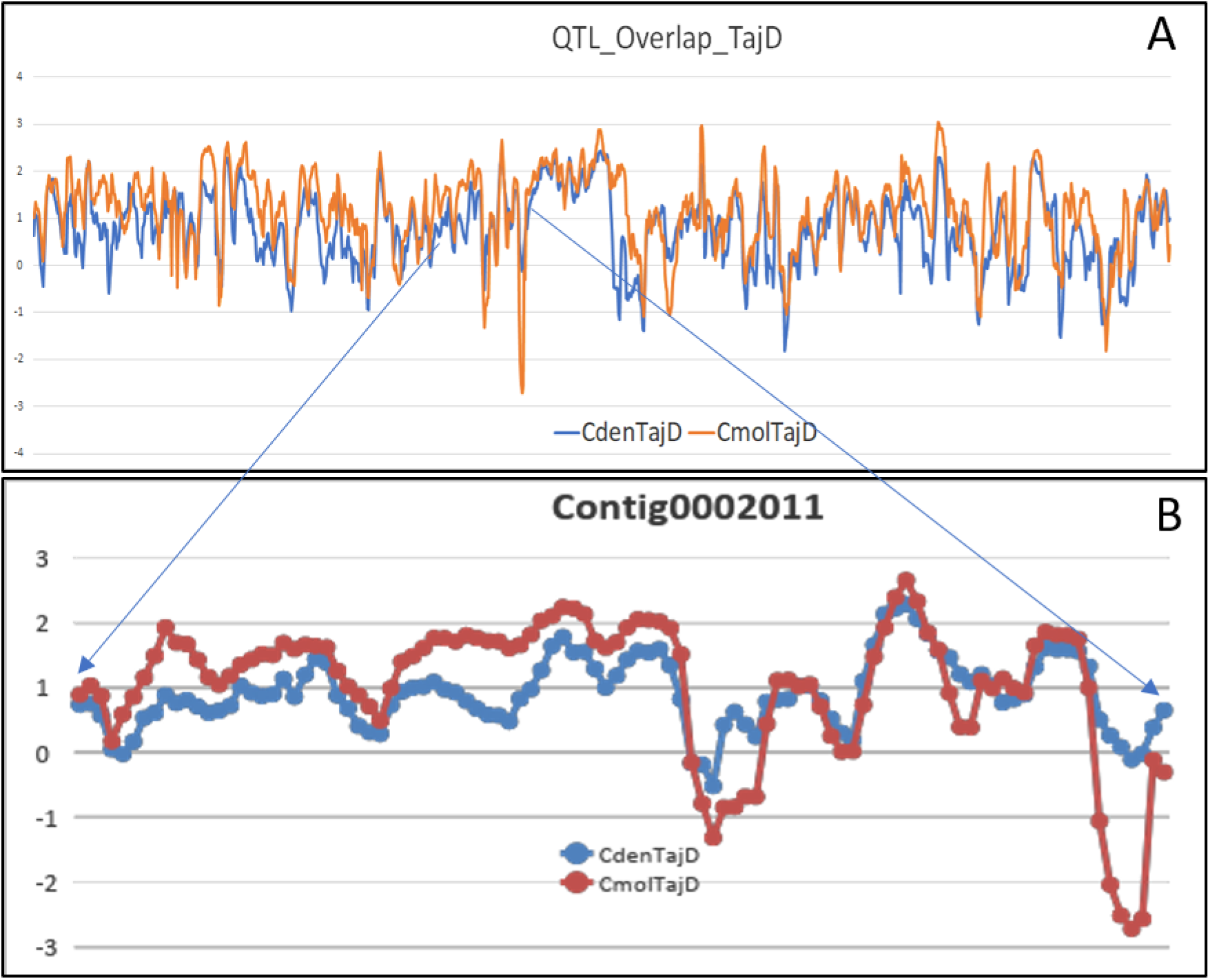
Stepwise signature of selection analyses of a region of overlap in the LGB QTL for blight resistance in two reference mapping studies. **A.** This is a profile of TajD values determined from nucleotide diversity ratios in the region of overlap of the QTLs region from the community reference genetic linkage map (9) and a higher density version (22), for cbr1 *Cryphonectria* resistance on LGB. The region of overlap consists of 13 contigs spanning a total of 1.122Mb. The strongest differential signature of selection (negative TajD value) in *C mollissima* is within one gene in contig 0002011 (se Fig XX below). **B.** This profile of TajD values was determined from nucleotide diversity ratios across the one contig (0002011) within the cbr1 *Cryphonectria* resistance QTL region on LGB. This contig maps to EST-DNA marker CmSI0550 on the reference genetic map (9). The *C. mollissima* profile suggests a very strong signature of selection at the end of this contig across a window of app. 6,000 bp.

<u>In parallel, we compared the datasets from the Asian species diversity analyses (see above)</u> to examine whether any of the 112 putative introgressed genes were located in any of three published QTL regions associated with resistance to chestnut blight (9). A total of 7 genes (Table 4) were found in these QTL regions, suggesting that the genetic basis underlying the blight resistance within the Chinese chestnut might be sourced from hybridization with the related species *C. seguinii*, which exhibits the highest blight resistance in our demographic analysis. Of these 7 candidate genes, 2 were located on Linkage Group B (see Supplementary Table S6 for detailed information on their locations on LGB). One candidate (maker-scaffold00115-augustus-gene-0.47-mRNA-1) was aligned by BLASTn to 6 locations on LGB, outside of the core region of the blight resistance QTL cbr1. However, the other candidate gene (gene model snap_masked-scaffold01565-abinit-gene-0.16-mRNA-1), aligned to a single position within the core of the major blight resistance QTL cbr1. This gene is highly similar in sequence to maf-like proteins (Mads Affecting Flowering2), which play a role in chilling requirements for bud break, and have been identified in several tree species (e.g. *Juglans regia*, Table 4).

**Table 4.** Seven genes identified by diversity signatures in the 3 blight-resistance QTL that may be candidates for introgression from *C. seguinii* into *C. mollissima*.

| Gene model and location | Strongest NCBI Blastn alignments |
| --- | --- |
| maker-scaffold00521-augustus-gene-0.34-mRNA-1 | *Quercus suber* protein LAZ1 homolog 1-like (LOC111989524), transcript variant X2, Evalue 6e-80, 97.28% identity, Accession XM_024021319.1) |
| augustus_masked-scaffold00172-abinit-gene-1.14-mRNA-1 | *Quercus suber* pentatricopeptide repeat-containing protein At1g02370, mitochondrial (LOC112016111), Evalue 0.0, 96.46% identity, Accession XM_024048621.1) |
| maker-scaffold00115-augustus-gene-0.47-mRNA-1 | *Quercus suber* uncharacterized locus (LOC112039279), transcript variant, ncRNA, Evalue 6e-134, 92.33% identity, Accession XR_002885976.1; |
|  | *Quercus suber* 50S ribosomal protein L11, chloroplastic-like (LOC111991637), Evalue 4e-85, 90.16% identity, Accession XR_002880753.1) |
| maker-scaffold10970-augustus-gene-0.7-mRNA-2 | short (~450 bp) exon-like matches to Juglans regia RNA-binding protein 39 (LOC108993348), Evalue 3e-143, 79.30% identity, Accession XR_001996491.1 |
|  | Quercus suber serine/threonine-protein kinase Kist-like (LOC112023857), Evalue 2e-127, 91.44% identity, Accession XR_002884076.1) |
| maker-scaffold10970-augustus-gene-0.7-mRNA-1 | short (~450 bp) exon-like matches to Quercus suber uncharacterized locus LOC112006296, partial mRNA, Evalue 0.0, 97.99% identity, Accession# XM_024038578.1 |
|  | Juglans regia RNA-binding protein 39 (LOC108993348), transcript variant, Evalue 3e-143, 79.30% identity, Accession# XR_001996491.1) |
| snap_masked-scaffold01565-abinit-gene-0.16-mRNA-1 | Juglans regia maf-like protein DDB_G0281937 (LOC108980279), transcript variant, Evalue 5e-63, 77.81% identity, Accession # XR_001994378.1) |
| snap_masked-scaffold04470-abinit-gene-0.10-mRNA-1 | Quercus suber subtilisin-like protease SBT4.15 (LOC111986162), Evalue 8e-61, 85.51% identity, Accession #XM_024017783.1) |

**Table 5.** Identification of NBS-LRR genes and Cysteine-Rich Receptor-Like Kinase gene models in the Phytophthora cinnamomi (Pc) resistance QTL region in C. mollissima Linkage Group E.

| IPS ID | Family Name | Number of Genes per Chromosome* |  |  |  |  |  |  |  |  |  |  |  |  |
| --- | --- | --- | --- | --- | --- | --- | --- | --- | --- | --- | --- | --- | --- | --- |
|  |  | A | B | C | D | E | F | G | H | I | J | K | L | Total |
| PF01657 | Salt stress response/antifungal | 4 | 4 | 6 | 8 | 30 | 1 | 0 | 0 | 0 | 3 | 0 | 5 | 61 |
| PF00069 | Protein kinase domain | 86 | 46 | 33 | 33 | 57 | 36 | 31 | 53 | 39 | 34 | 33 | 34 | 515 |
| PTHR34630:SF3 | Disease Resistance Protein (TIR-NBS-LRR CLASS) Family | 0 | 0 | 0 | 0 | 1 | 0 | 0 | 0 | 1 | 0 | 0 | 1 | 3 |
| PTHR32099:SF34 | Cysteine-Rich Receptor-Like Protein Kinase 34-Related | 1 | 0 | 0 | 2 | 4 | 0 | 0 | 0 | 0 | 0 | 0 | 0 | 7 |

\* Pseudo-chromosomes named according to Linkage Group designations in chestnut research community reference genetic map (9).

#### Signatures of selection in the *Phytophthora cinnamomi* (Pc) resistance QTL region in *C. mollissima* vs *C. dentata*

Employing a similar analysis to that above, the distribution of transformed nucleotide diversity in American (*Pi Cden*/*Pi Cmol*) and Chinese (*PiCmiol/PiCden)* chestnut pools plotted along the *C. mollissima* v3.2 pseudochromosome 5 (LG_E) are shown in Fig. 9. Nucleotide diversity in *C. mollissima* pool aligned against the *C. mollissima* reference genome was lower than that in the *C. dentata* pool. In total 49 genomic regions (5kb and larger) were detected as candidate regions (CKR) under purifying selection in the *C. mollissima* pool.

**Figure 9.**
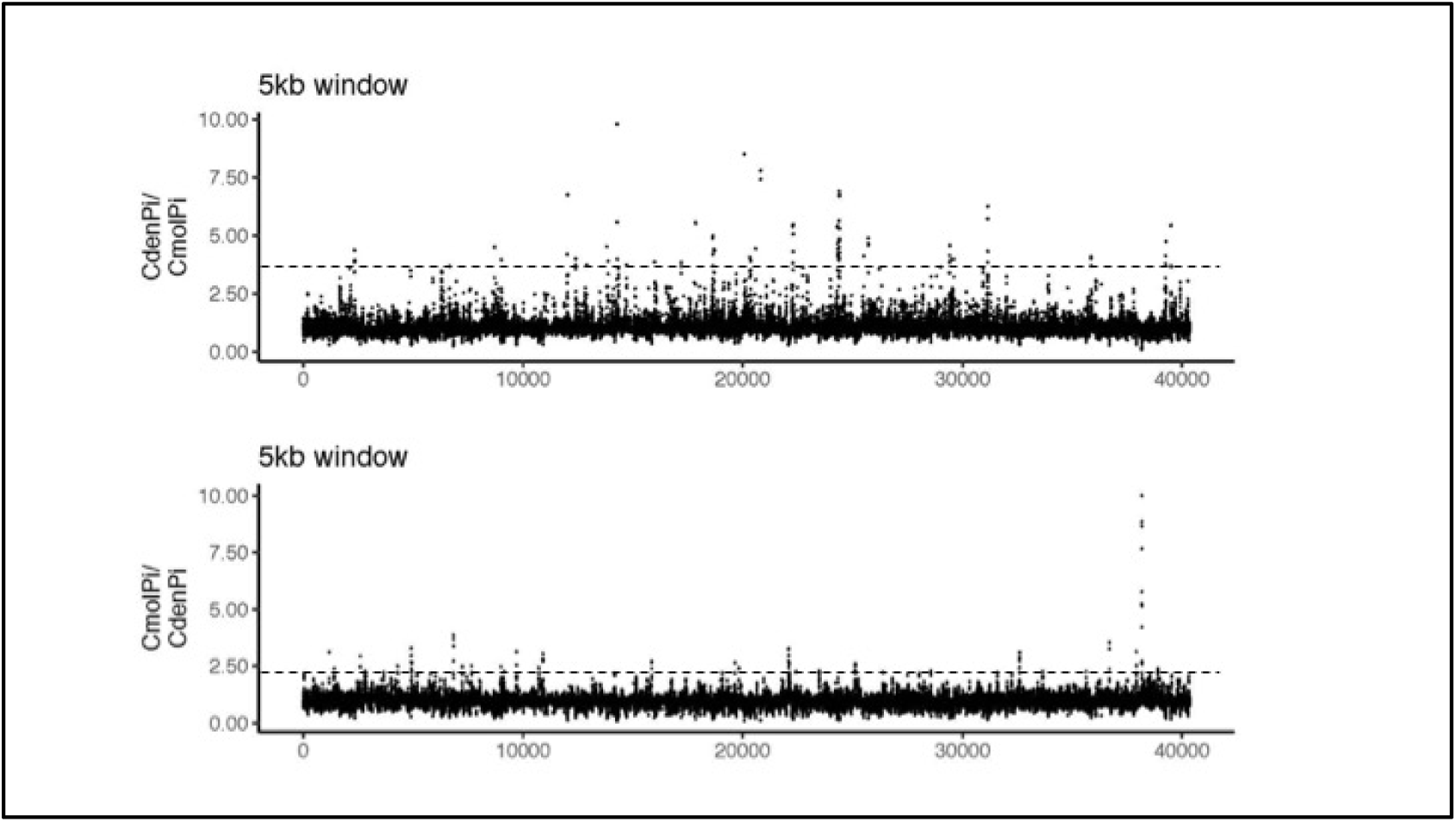
Distribution of transformed nucleotide diversity in American (*Pi Cden*/*Pi Cmol*) and Chinese (*PiCmiol/PiCden) chestnut* pools. Plotted along *C. mollissima* (Vanuxem) pseudochromosome 5 (LG_E). A dashed horizontal line indicates the cut-off used for identification of outliers representing the top 0.5% of nucleotide diversity ratios.

Leveraging data from previous QTL mapping studies of Pc resistance in Chinese/American hybrid families (23, 24), we ran statistical tests for neutrality across LG_E that we previously determined had three strong QTL intervals associated with Pc resistance. Using sequence-based markers from our mapping analyses and local blast alignment tools, we delineated the QTL intervals on the assembled Chinese chestnut LG_E pseudochromosome and determined if the statistical tests for neutrality detected any significant selection signatures in these QTL regions. Of the 49 regions of LG_E that exhibited purifying selection signatures in *C. mollissima*, 34 candidate regions were located within these QTL intervals (Supplementary Table S7 and Fig. 10). Similarly, 45 regions exhibited signatures of purifying selection in the *C. dentata* pool, but these are located outside of QTL intervals for resistance to *P. cinnamomi*. Most of the loci potentially contributing to adaptation Chinese chestnut to biotic stress caused by *P. cinnamomi* belong to genes involved in cell-wall formation, transmembrane signaling and transport, posttranslational protein modification and formation reactive-oxygen species (ROS) (Table S7).

**Figure 10.**
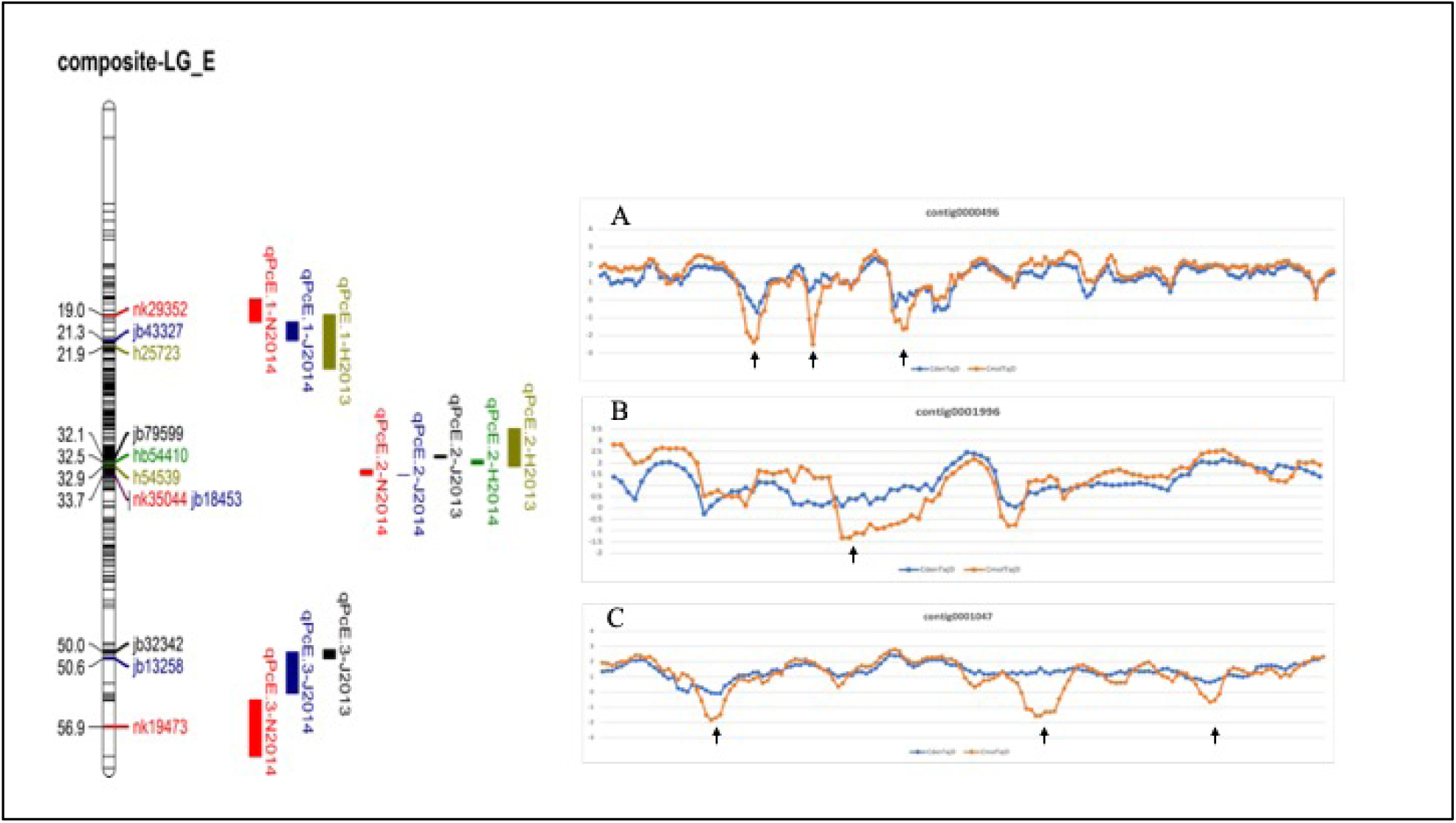
Three QTL intervals for resistance to *P. cinnamomi* in Linkage Group E. The TajD profiles for the 3 QTL regions (profiles A, B, C, respectively, for QTL qPcE.1, qPcE.2, qPcE.3) are shown on the right. TajD peaks of approximately −2 were considered most significant (identified with arrows), in which candidate genes for disease resistance were identified. The relative positions of selected candidate genes and the location of DNA marker loci associated with these genes are shown with the genetic linkage map on the left.

#### Signatures of selection in phenology traits, the bud burst QTL region in Chinese chestnut, oak and peach

Due to the extensive colinearity of deciduous tree genomes as highlighted above, we were able to perform genome comparative analysis for mapped QTL controlling budbreak in peach (25), oak (Bodenes, C., INRA Pierroton, unpublished results), and Chinese/American chestnut hybrids (22). This analysis revealed one common major colocalizing QTL region that in all three mapping analyses contributed with highest significance to variation for budbreak either floral buds (peach) or vegetative buds (oak and chestnut). This QTL was originally mapped in peach (25) and corresponds the location of the DAM genes in Prunus (26). Here, we performed a comparative sequence characterization of this region among these three species utilizing the published genome sequences of peach v2 and oak (*Q. robu*r) and the Chinese chestnut version 3.2. (Fig. 11). Results from this sequence comparative analysis reveal a high degree of preserved gene content and order among these species genomes, however, the Fagaceae species do not contain the tandem duplication of DAM genes that is characteristic of the peach genome. Due to the high degree of gene preservation in this region across many species (Fig. 11), we hypothesized that genes in this region could show signatures of selection among chestnut species particularly since budbreak timing is a very selectable trait in fruit trees and Chinese and American chestnuts show significant differences in the budbreak dates (22, Hebard, F.V., personal communication). In order to determine if any of the genes in this common budbreak QTL demonstrated signatures of selection and if so which genes, we performed a Tajima’s D analyses for LGL in chestnut as outlined for the studies we performed on LGE and LGB, above. The analysis showed that across linkage group L, 43 loci were identified with negative TajD values in *C. mollissima* and with neutral or positive values in C. *dentata*, indicating regions of differential selection (purifying selection). However within the QTL region, only the DAM gene orthologue in chestnut showed a signature of purifying selection in the chestnut species comparison (Figure 11). This is consistent with the importance of its putative role in controlling dormancy and bud flush and the high heritability and selectability of this trait in fruiting trees.

**Figure 11:**
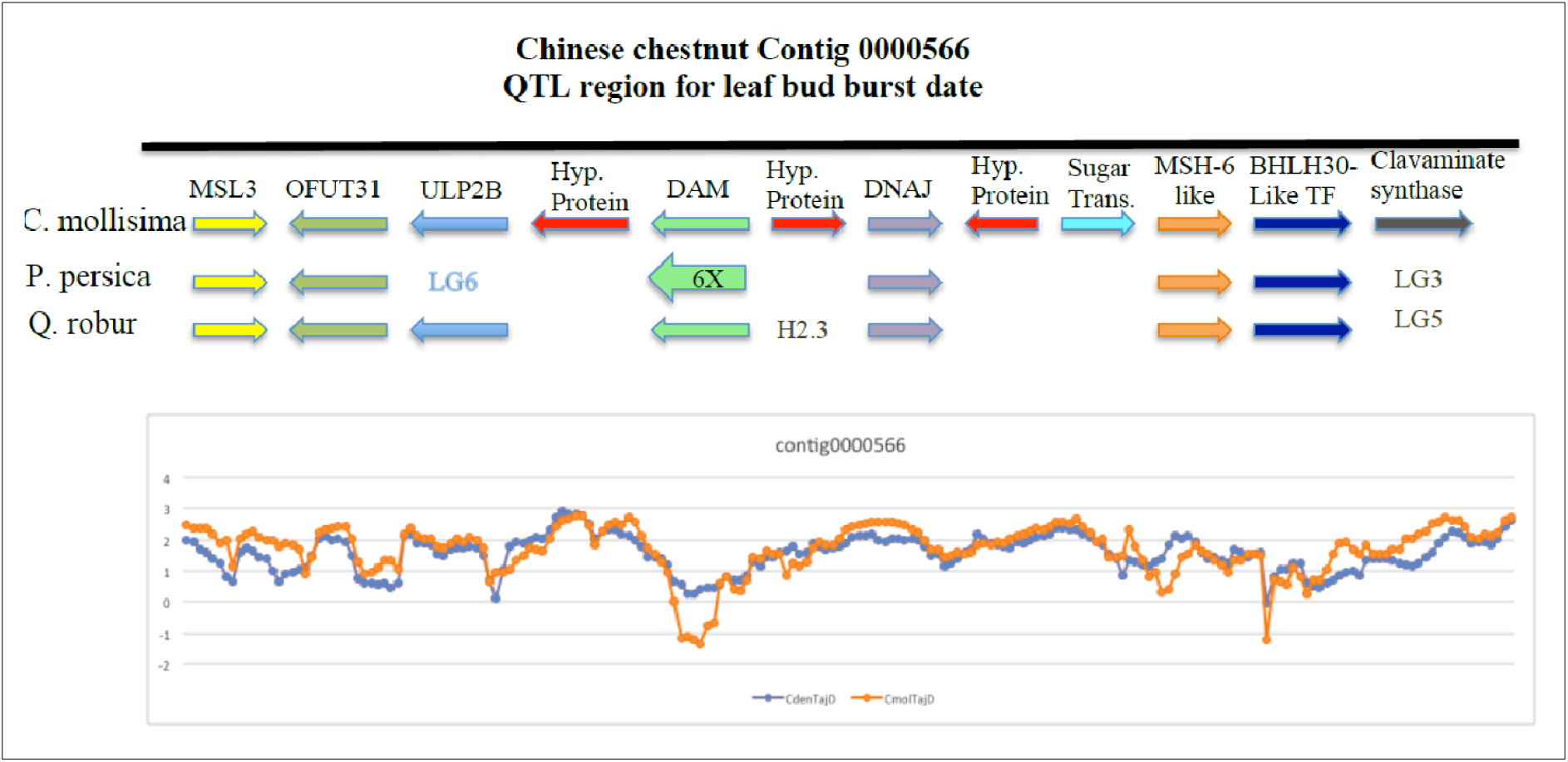
The structural organization of the DAM gene-containing region located in mapped QTL intervals of vegetative bud break in chestnut and oak and floral budbreak in peach. Arrows denote specific annotated genes of the three species showing colinearity. Where arrows are not present denotes the gene is not found in this region and written designations give the alternative location if it is annotated on the respective genome. 6X refers to the Prunus specific 6X segmental duplication of the DAM genes. The lower plot depicts the TajD analyses of this region demonstrating the differential purifying signature of selection in the *C. mollissima* genome (orange line) vs the *C. dentata* genome (blue line).

## DISCUSSION

The tragic story of the American chestnut’s demise is just one sobering example of what is becoming a recurring theme of the unintentional impact of human activity on our natural forest ecosystems. The success of future forest conservation and restoration efforts will increasingly rely on our ability to rapidly generate genetically improved tree materials for forest replanting. Due to the long generation times for many of our heritage forest tree species, traditional tree breeding approaches on their own do not offer an adequate solution to meet this challenge as woefully illustrated by the complete loss in ∼thirty years of the American chestnut as a dominant species in eastern North American forests in the early part of the 20^th^ century. Traditional backcross breeding methods to introgress resistance to *C. parasitica* from Chinese chestnut into American chestnut over the last ∼30+ years have only achieved limited success. However, the developing tools of genomics science coupled with traditional breeding practice and biotechnical approaches afford the opportunity to more rapidly develop improved genetic materials to meet the environmental challenges imposed by human activity. As the foundation of a genomics toolbox, a whole chromosome scale genome sequence for a species provides the genetic architecture by which we can bridge genetic studies of traits to discovery of the underlying genes that control these traits. With this goal in mind, we developed a chromosome scale whole genome sequence assembly of the Chinese chestnut genome and implemented it for evolutionary and comparative genomics studies to advance our understanding of genome evolution and adaptation of species in the genus Castanea and to identify candidate genes for traits critically important to future conservation and restoration efforts.

### The Chinese chestnut genome assembly and structural features

As shown in Supplementary Table S1, the development of a chromosome-scale genome for the *C. mollissima* cultivar Vanuxem, required a long, step-wise manner approach through several *de novo* assemblies, gap closings, and anchoring of scaffolds to the chestnut research community’s reference genetic linkage map. The initial draft *de novo* assembly of the genome, version V1.1, which was released to the public in January 2014 (www.hardwoodgenomics.org), consisted of 41,260 scaffolds covering 724 Mb (an estimated 91.2% of the Chinese chestnut genome). The V1.1 genome draft has supported many investigations and publications (27, 28, 29, 31). To better support basic research and restoration of the American chestnut, we then focused efforts for several years on developing contiguous, chromosome-scale sequences. This proceeded through painstaking manual merging of contigs and gap-closing until an assembly (V3.2) of 12,684 contig sequences spanning 783.4 Mb was achieved. Assembly of draft pseudochromosome sequences (version V4.0) was accomplished by anchoring 4,314 of the V3.2 contigs to the chestnut research community’s reference genetic linkage map (9). The overall placement of contigs was validated through cytological mapping (Fig. 2 and Fig. 3), graphical visualization of linkage group loci and pseudochromosome sequence matches (Fig. 1), and chromosome-scale sequence alignments to other tree genomes (Fig. 4 and Fig. 5), and. However, our scaffolding and anchoring approach left gaps of unknown size and content between each contig, and the total pseudochromosome assembly represented only 58% of the V3.2 *de novo* genome assembly of 783Mb. By conservatively removing markers that identified more than one contig in the pseudochromosome assembly, we may have biased our pseudochromosome builds for gene rich regions over repetitive non-coding regions. In addition, the availability of only short read sequences during the *de novo* stage of contig assembly would also have limited the extent of repetitive DNA assembly. Nevertheless, the Circos plots of single pseudochromosome to genome comparisons (figures S1 and S2) did reveal the presence of some dispersed repeats in the V4.0 chestnut genome assembly. Future improvement the assembly may be achieved through the use of more recent long-read technologies, such as Nanopore (32), and scaffolding with chromatin-interaction data, such as Hi-C (33) However even these approaches may result in less than full-genome assembly given the challenges of high heterozygosity levels and an inability to generate dihaploid individuals in Chinese chestnut. The recently published *Quercus robur* genome (13) utilized synteny with the *Prunus persica* genome to order incorporate contigs and scaffolds that had not been assembled *de novo* nor scaffolded with oak genetic map markers. This approach assumes that micro-level syntenies follow known macro-syntenies based on genetic maps, which may not always hold true. However, the hybrid synteny approach could also complement long-read technologies in future Chinese chestnut genome improvements.

The current V4.0 assembly, although an incomplete draft, does by virtue of chromosome-scale sequences provide a significant advancement in our ability to investigate genome organization and the evolution and genetic structure of important traits such as disease resistance, as well as applications such as genome-wide selection. Our comparisons of the chestnut genome with a selection of genomes that include the herbaceous model plant Arabidopsis, as well as woody vines and trees (grape, oak, peach, and poplar) confirmed that there have not been any recent genome duplications, in keeping with the reports for peach and oak. Our comparative analyses also confirmed the strong colinearity among genomes and gene content in tree species (29). This conservation of genome structure and information content after millions of years of species divergence suggests strong constraints on the evolution imposed in perennial plants of long-generation time and limited domestication. The conservation of genome structure may also be an underlying reason that gene flow is high and that inter-species natural hybridizations in natural stands of trees are so common. The conservation of genome structure and information content will permit the leveraging of information from model plants and among tree species to more rapidly advance our understanding of the many unique and fascinating features of long-lived tree species.

### Genome diversity and selection for disease resistance in Asian chestnut species

Before the last glacial period, the effective population sizes of the three Asian Castanea species *C. seguinii*, *C. mollissima* and *C. henryi*, declined in fluctuations, which corresponded to the multiple and cyclical changes of climate during the Quaternary period. During the last glaciation period, the three species congruently experienced a similar bottleneck, followed by a period of rapid and asynchronous growth starting from the last glaciation maximum (LGM), suggesting that climate change played important role in shaping the demography of the Chinese chestnut. The effective population sizes peaked around 4,000 - 10,000 years ago for three species. Then, similar to the population shrink in American chestnut, the Ne of *C. henryi* reduced steeply during the China’s slavery and feudalism, but the Ne of *C. seguinii* was nearly stable. These asynchronous changes of Ne-s reflecting that *C. seguinii* bore the highest blight resistance and the blight resistance in *C. henryi* was weaker than that in *C. mollissima* and *C. seguinii*. Given that the chestnut blight was a deadly epidemic for *C. henryi* trees, demography reflected that the initial infection of *Cryphonectria parasitica* on chestnut trees in China would have occurred in the most recent 4,000 years. An alternative explanation to the asynchronous decreases of effective population sizes is the effects of anthropogenic activities since the agrarian age, especially the changes of land use in agriculture. However, the three *Castanea* species always distributed adjacently in China and bore differentiated resistance to the chestnut blight, we inferred that their asynchronous changes of Ne-s recently may be caused by their different tolerance for the common diseases. *Furthermore, their different incidence levels and demographics indicate that more heritable resistance to chestnut blight may have been recently accumulated in genomes of C. mollissima and C. seguinii than in C. henryi*.

This evolutionary relationship indicates a possible single origin of high blight resistance within the *C. mollissima* and *C. seguinii* populations, but their divergence time was dated at around 484.57 thousand years ago, which was much earlier than the inferred upper boundary of initial infection time from demography above. The effective population sizes of the three Asian *Castanea* species after their divergence decreased to the lowest level during the last glacial period but rebounded from the LGM period. Therefore, we infer that the blight resistance was not inherited from their common ancestor, at least not completely inherited.

In summary, our population genomic analyses showed that *C. seguinii* responded to the blight fungus rapidly with minimum loss of genetic diversity in the past hundreds of generations, which facilitated the adaptation of Chinese chestnut trees through hybridization and introgression in their common distributional regions. At large time scales, climate change during the Pleistocene, especially the last glacial period, may have influenced the genomes of these three Castanea species as well.

### Signatures of selection for blight resistance on Linkage Group B in *C. mollissima* vs *C. dentate*

Results from analyses of the Asian chestnut species (above) suggest that pathogen selection pressure and interspecies hybridization from one species to another may underpin the resistance to *C. parasitica* in *C. mollissima*. Under such a hypothesis American chestnut would not be expected to show signatures of selection on the same genes as Chinese chestnut in genetically mapped resistance QTL intervals. Resistance to *C. parasitica* has been mapped in several interspecies hybrid Chinese x American families in both F2 and backcross configurations (9, 22). From this analysis it appears that variation in the resistance phenotype is attributable to a relatively small number of loci donated by the Chinese parent in the initial cross and that one region in particular, located on linkage group B is reproducibly associated with the resistance. For this reason, leveraging the *C. mollissima* whole genome sequence, we performed a comparative Tajima D analysis across linkage group B in five *C. mollissima* and five *C. dentata* resequenced accessions and searched for contrasting evidence of selective sweeps between the species genomes. A stepwise signature of selection analysis of a region of overlap between the LGB blight-resistance QTLs region from the community reference genetic linkage map (9) and a higher density map (22) resulted in the identification of only one gene in the overlapping QTL region with a highly significant signature of purifying selection in Chinese chestnut but not American chestnut. This gene model is a putative inositol polyphosphate-related phosphatase gene. Inositol phosphate signaling has been linked to multiple effects within plants and most notably for our study abiotic stress (34) and plant defense responses (35, 36). Thus, this is a strong candidate gene and our selective sweep results are consistent with several hypotheses. One hypothesis would suggest that the resistance in *C. mollissima* derives from a founder effect of an introgressed genome segment from a resistant *C. seguinii* and this resistance was selected during coevolution of the host and the pathogen as suggested above. Another hypothesis is that this signature in the *C. mollissima* genome is a direct result of selective forces imposed by the co-evolution of the *C. mollissima* genome under the pathogen selection. In either case, we would not expect evidence of purifying selection on this gene in *C. dentata* since historically, as far as we know, the pathogen was not present in North America prior to its introduction in the turn of the 20^th^ century. Finally, as we can perform transgenic studies in American chestnut, this candidate can be directly tested for its ability to confer resistance in future transgenic experiments.

### Signatures of selection for Phytophthora cinnamomi resistance in C. mollissima vs C. dentate

*Castanea* species originated in eastern Asia, moved westward during the Tertiary period and currently exhibit a disjunct distribution pattern in eastern Asia and eastern North America (37). Based on the global studies of mating type, *P. cinnamomi* is also hypothesized to have an Asiatic origin (38, 39). Thus, co-evolution between chestnut and this pathogen in Eastern Asia could have generated a strong selection pressure to evolve defense mechanisms directly targeting *P. cinnamomi* and/or blocking host infection. Nonrandom mutation rates within genomic regions under selective pressure (i.e. selective sweep) can be detected using variety of population statistics (e.g.. neutrality tests estimating nucleotide diversity). Because of the dominant inheritance of *P. cinnamomi* resistance (40) and high infectivity of geographically different isolates, purifying (or negative) selection on genes potentially involved in resistance could play a significant role in the adaptation of Eastern Asian chestnuts to stress caused by *P. cinnamomi*. This mechanism would reduce the genetic diversity of these genes by elimination of susceptible alleles from the population followed by a random mutation process. Excess of rare mutations and negative TajD values could be signatures for genomic regions that passed through this bottleneck of purifying selection.

The plant cell wall is the first barrier encountered by *P. cinnamomi* zoospores attempting to colonize chestnut roots. Alterations in cell wall structure have significant impact on disease resistance (41, 42). As indicated by the neutrality test, several cell wall-associated genes within QTL intervals were under purifying selection in the *C. mollissima* but not in the *C. dentata* pool. These are a probable pectin methylesterase CGR2 (contig0000773_177500 - 180500) and a Golgi-localized type II membrane protein, that has enzymatic activity toward pectin methyl-esterification. Two genes involved in phenylpropanoid metabolism, a probable 4-coumarate-- CoA ligase 1 (contig0003825_11500 - 13500) and a flavanone 3-hydroxylase (contig0001240_19500 – 22500) may contribute to lignin biosynthesis and modification.

The β-linked glucose polysaccharides are the most abundant component of *Phytophthora* cell walls. They present a very complex array of possible structures, some with well-established activity in modulating plant innate immunity (43,44). In resistant avocado rootstocks β-1,3-glucanase and callose inhibit zoospore germination and subsequent hyphal growth of *P. cinnamomi* (45). Also, glucan endo-1,3-beta-glucosidases were overexpressed in *C. crenata* roots infected with *P. cinnamomi* zoospores (46). In our neutrality test, a putative endo-1,3-beta-glucosidase (contig0000723_100500 – 106500), a member of glycoside hydrolase family 17, was under selective sweep in *C. mollissima*.

A putative beta-fructofuranosidase, an ortholog of insoluble CELL WALL INVERTASE 1 (CWINV1) in Arabidopsis was under selective sweep within the QTL1 region (contig0000278_33500 - 36500). This enzyme is ionically bound to the cell wall and was described as one of key enzymes during plant pathogen/interactions. The degradation of apoplastic sucrose by CWIN is crucial for biotrophic and hemibiotrophic pathogens as they mainly rely on the sugar acquisition from the apoplast through the activity of hexose transporters (47, 48).

Membrane-localized receptors are the second line of defense against pathogens. Chemical substances associated with pathogen and/or cell wall degradation can modulate plant innate immune response upon recognition by receptors with varied extracellular domains. Two receptor-like genes, the G-type lectin S-receptor-like serine/threonine-protein kinase G-LecRK (contig0000723_29500-34500) and a block of duplicated cysteine-rich receptor-like protein kinases CRKs (contig0001047), were under purifying selection in *C. mollissima*. The G-LecRKs G-type lectin S-receptor-like serine/threonine-protein kinase confers resistance to biotic stresses and fungal pathogens in Arabidopsis and other crops (49). They have an extracellular sugar-binding domain that may perceive the oomycete-associated chemical signals to trigger innate plant immunity (44). The G-type-lectin-RLKs were upregulated in roots of diploid strawberry, citrus rootstocks and Japanese chestnut infected with *P. cactorum*, *P. parasitica* and *P. cinnamomi* zoospores, respectively (50, 51, 52).

The CRKs are transmembrane proteins that exhibit ectodomains containing the cysteine-rich Domains of Unknown Function 26 (DUF26). They constitute a land plant-specific family of carbohydrate-binding proteins expanded through tandem duplications. A block of tandem duplicated CRKs within qPcE.3 was identified as having a potential selective sweep (with negative TajD values) in Chinese chestnut (Fig 8). Due to presence extracellular Cys-rich domains (C-X8-C-X2-C), the CRKs are potential targets for redox modifications and hypersensitive response associated with programmed cell death (53, 54). They act in non-redundant fashion in response to biotic stress difference (55, 56). The CRKs were overexpressed in soybean roots induced by *P. sojae* zoospores (57) and in resistant *C. crenata* genotypes treated with *P. cinnamomi* zoospores (52). Noteworthy, homologs of the protein disulfide isomerase-like 1-5 enzyme (contig0000496_63500 - 66500) with a protein disulfide isomerase activity catalyzing the rearrangement of the -S-S- bonds in proteins, was also under selective sweep in the *C. mollissima* pool as wells as a tandemly duplicated ABC transporter C family member 8 (contig0000423_120500 – 144500) which functions as a pump for glutathione S-conjugates in transmembrane transport (58).

Activating mechanisms of innate plant immunity by reprogramming host cell molecular network is the third line of defense against pathogen spread. Two intertwined posttranslational modifications, protein ubiquitination and phosphorylation, play essential roles in intracellular signal transduction triggered by plasma membrane-resident receptors. In coevolution of the *C. mollissima- P. cinnamomi* host-pathogen system, Putative ubiquitin protein ligase 5 (contig0000302_26500 – 31500) and mitogen-activated protein kinase kinase kinase 17 (contig0003275_32500 – 37500) may represent elements of defensive networks shared with receptor-like kinases.

The most striking difference in nucleotide diversity between *C. dentata* and *C. mollissima* was observed in the middle of the chr E colocalizing with qPcE.2, the most stable QTL detected multiple years progeny derived from two Chinese chestnut sources of resistance to *P. cinnamomi*. Two genes were annotated in this region – a putative ALA-interacting subunit 2, homolog of the ligand-effect modulator 3 (AT5G46150) in Arabidopsis that encodes a protein of unknown function with transmembrane activity; and an ortholog of ornithine delta-aminotransferase (KEGG:AT5G46180) involved in proline metabolism and transcriptionally up-regulated in response to osmotic stress and non-host disease resistance (59). At the molecular level, proline accumulation may act as antioxidative defense system by directly scavenging free radicals and preventing programmed cell death (60). On the other hand, proline may act as osmolyte mitigating water stress and balancing turgor pressure during abiotic stress and pathogen invasion at a whole-plant level (61,62). Wilting caused by blockage of xylem by the pathogen and reduction of water supply, is a classical symptom of a root disease on infected chestnut plants (63). Finally, proline may act directly on pathogen propagation and spread because of its significant role in osmoregulation of development and release the *P. cinnamomi* zoospores (64). Thus, neutrality tests using the whole-genome resequencing datasets *C. mollissima* and *C. dentata* highlighted potential importance of ornithine delta-aminotransferase for adaptation *C. mollissima* to pathogenic pressure by *P. cinnamomi*.

### Castanea comparative genomic analyses

In this study we conducted comparative analyses of the genomes of chestnut and oak, both species of the Fagaceae, and with peach, a member of the Rosaceae. From these comparisons we infer that, as seen in oak, there have not been any whole genome duplication events in chestnut after the evolutionary splits of the *Quercus* and *Castanea* genera, and of the Fagaceae and Rosaceae families. Additionally, the high level of preservation of genome organization among these tree species again further underscores the hypothesis that deciduous tree genomes could be more slowly evolving than genomes for plants with different life traits (29, 65, 66).

In hardwood forest trees, traditional single family trait mapping and selections typically have not been done due to the long generation times of these species. In contrast, fruit tree genetics that is driven by domestication and orchard plantings is a rich resource of genetic information on genes that control many aspects of fruit tree growth and development (18). As fruit trees are deciduous, many trait/gene associations are likely to translate to hardwood forest trees. Here we demonstrate that the high level of genome synteny between the peach, chestnut, and oak enables a comparative QTL analyses for traits central to sustaining forest trees in a rapidly changing environmental landscape. Our initial results indicate that trait/gene associations in one tree species may easily be translated to others providing key information for genetic improvement of tree species with less genomic resources.

### Signatures of selection for bud burst QTL region in chestnut, oak and peach

Arguably from the standpoint of rapid climate change, the rate of evolution of tree genetic composition and the rapidly changing environmental factors pose one of the greatest challenges to adaptation of perennial trees particularly for phenological traits such as flowering and vegetative bud burst. From a number of studies in fruit and forest trees, a picture of the genetic control and evolution of the genes and gene networks that control the timing of floral and vegetative bud break is emerging (67,68,69). Comparative mapping of QTL locations of budbreak loci among peach, chestnut and oak and the availability of whole genome sequences for each species enabled us to quickly surmise that an orthologous genomic region of all three species was present in the major budbreak QTL of each species. This region contains a single MADs-box transcription factor gene in oak and chestnut and a segmentally duplicated gene (six copies in Prunus) that has previously been characterized as a major floral bud dormancy and budbreak control gene in a number of fruiting tree species (26, 69, 70) and in at least one fruit tree vegetative budbreak QTL as well (71). Combining comparative genomic sequence analyses, comparative QTL analyses and our TajD analyses of linkage group L in chestnut, we hypothesize that the DAM gene containing locus in these deciduous forest tree species is a major control locus for both vegetative and floral budbreak and in the case of the DAM gene due to its central importance in regulating the timing of budbreak as seen in fruiting trees, differentially predisposes it to environmental and breeding selection pressures over those other genes in this conserved region.

## Data availability

The contig and scaffold sequences have been submitted to NCBI, and are also available for download and query at the Hardwood Genomics Project website (https://www.hardwoodgenomics.org/genomes).

## Acknowledgments

This project was funded by The Forest Health Initiative through grant # 137RFP#2008-011 to JEC. Support was also provided by the USDA National Institute of Food and Agriculture grant 2016-67013-24581 to the American Chestnut Foundation. Additional support was provided through several grants-in-aid to JEC from The American Chestnut Foundation and to JEC through the USDA National Institute of Food and Agriculture Federal Appropriations under Project PEN04532 and Accession number 1000326.

## MATERIALS AND METHODS

### A.1. Tree materials and Sample preparation

#### Reference tree

Leaves and twig tissues were collected from the Chinese chestnut blight-resistant *Castanea mollissima* genotype ‘Vanuxem’ (Supplementary Table S8) at The American Chestnut Foundation’s farm in Meadowview, VA. Collections in summer 2011 were used for the version 1.1 gDNA assembly, while tissues collected in the summers of 2014 and 2016 were collected for version 2 and 3 assemblies. The Vanuxem genotype was chosen for sequencing because it is expected to remain readily available to breeders and researchers. Also the Vanuxem genotype was used as a resistant parent in crosses for published genetic linkage maps (1) and as the source DNA for the BAC libraries used in constructing the physical map and integrated genetic-physical map for Chinese chestnut (2). The cultivar Vanuxem was also chosen as the least heterozygous (50%) among several Chinese chestnut blight-resistance gene donor parent trees (ranging from 52% to 64% observed heterozygosity) within The American Chestnut Foundation’s breeding program, as determined with 25 Simple Sequence Repeat loci. Tissue samples were immediately snap-frozen in liquid nitrogen and then stored at −80°C. DNA was extracted for 454 and Illumina genome sequencing from bud, cambial, and leaf tissues using a modified CTAB protocol (3). DNA was extracted for PACBio sequencing by the Arizona Genomics Institute from 36-hour dark treated (tarp-shaded) leaf samples.

#### Diversity Panel for QTL analyses

Twig and leaf samples were collected, and immediately snap-frozen in liquid nitrogen, in early spring of 2015, from five *C. dentata* genotypes and *six C. mollissima* genotypes (Supplementary Table S8). Tissue samples were provided by the Connecticut Agricultural Experiment Station (CAES), by The American Chestnut Foundation (TACF), and by the Pennsylvania Chapter of The American Chestnut Foundation (PENN). DNA was extracted for Illumina HiSeq library construction from the twig and/or leaf tissues using a modified CTAB protocol (3).

#### Genome Diversity and Evolution Panel for Asian species analyses

For diversity analyses of the Asian chestnut (*C. mollissima*, *C. seguinii* and *C. henryi*) species genomes, 98 trees across 76 locations spanning the geographic ranges of the three species were sampled (Fig. 6, Supplementary Table S8). To deduce the effects of domesticated Chinese chestnut trees, we filtered out populations <50 km far from human settlements and man-made chestnut forest. In each population, sampled trees were at least 500 m apart. Fresh leaves were collected from first year branches and silica gel dried leaves were used in DNA-seq.

### A.2. Genome sequencing and assembly

#### Genomic DNA sequence

Over 61 billion bases of genomic DNA sequence data were produced from a combination of Illumina MiSeq and Roche 454 Next Generation Sequencing platforms. This included twenty-one 454 FLX sequencing runs on 454 FLX machines, producing 25,179,431 reads averaging 516 bp in length and totaling 13,175,668,630 bp of sequence. Also 915,895,342 bp of BAC-end sequences were obtained by 454 FLX paired-end sequencing of pools of BAC clones tiling the physical map of the Vanuxem cultivar (2) to 1.5X depth. In addition, 9 runs of 250 bp paired-end reads of a 480 bp insert Illumina genomic DNA library on MiSeq machines produced 41,300,000,000 bp of sequence. The Chestnut physical map (1) minimum tiling path of BAC clones were also sequenced in 2 runs of 250 bp paired-end reads on the MiSeq, producing another 4,700,000,000 bp of sequence. Finally, two long insert libraries averaging 3,000 bp and 8,000 bp were prepared for 454 FLX sequencing, yielding 897,238 and 884,030 mate-pair reads averaging 500 bp per read, totaling 890,634,000 bp of sequence for use in scaffolding. Overall, the 454 FLX and MiSeq data totaled 60,982,197,972 bp of high-quality sequence data, representing app. 76X depth of coverage of the 794 Mb genome (4).

#### Draft Genome Assembly and Scaffolding

Ten hybrid assembly builds using the Newbler assembler versions 2.5, 2.6, and 2.8 were conducted with various amounts and combinations of 454 and Miseq data. The best hybrid assembly was obtained from the 7^th^ assembly, using the heterozygosity option in Newbler v2.8. The total number of input reads was 89,135,536 (covering 36,739,712,156 bp), of which 77,421,025 reads were included in the final assembly. The assembly included input of 9,096,315 Illumina MiSeq paired reads (of which 5,192,637 paired reads were assembled into the same scaffolds at an average distance of 566 bp); along with a total input of 897,238 paired reads from the 454 3 kb insert library (of which 529,560 paired reads assembled in the same scaffolds at an average distance of 1,804 bp); and a total input of 884,030 454 8 kb insert paired reads (of which 507,004 paired reads assembled in the same scaffolds at an average distance of 6,076 bp).

#### Final *de novo* Genome Assembly

The de novo genome assembly was improved through gap-closing and contig merging prior to building chromosome-length sequences. For this, 6.8 Gb PACBio sequence data was generated by the Arizona Genomics Institute from flash-frozen etiolated leaves collected directly from the Vanuxem ramet at the American Chestnut Foundation’s farm in Meadowview, VA, that was previously sampled for 454 and Illumina sequencing. Filtering of low quality reads, removal of short reads, and sequence correction using Illumina reads and the de novo contig sequences, yielded 2 Gb of high quality long sequence reads. The PacBio reads were error corrected using the CLC PacBio Correction tool. Corrected PacBio reads were pooled along with high-quality consensus sequences generated from mapping 454 and MiSeq reads against a set of assembly contigs that were already in the process of editing and clean-up. Mapping and consensus generation were done using the CLC Read-Mapping and Consensus Sequence tools. The pooled PacBio reads and consensus contigs were meta-assembled using the overlap layout consensus algorithm of the De Novo Assemble tool in Genious. This meta-assembly produced ∼72k contigs greater than 1 Kb with a max of 594 kpb, spanning 812 Mb. The meta-assembly in Genious was followed by multiple rounds of contig-joining/gap-filling using the long-read algorithm of the Join Contigs tool in CLC Genomics Workbench. Successive rounds of contig joining were performed, where each round used as input a set of long-reads generated by re-mapping MiSeq and 454 reads against the new consensus contigs, saving un-mapped reads, and using those as input for PacBio read correction. In this way the “gaps” in the assembly were closed by segments of PacBio reads corrected by unmapped (i.e. gap-spanning) short 454 and MiSeq reads. Joined contigs were then ‘cleaned’ by mapping a comprehensive set of Vanuxem short read data (454SE, 454MP, MiSeq, whole-genome and transcriptome data) and generating consensus sequences using a majority-rule condition to minimize ambiguities in the final sequences. Contigs were split at points where total read depth was low (<5X) and only consensus sequences over 5 Kb were kept. Gap filling and contig merging with the PACBio reads produced longer, more contiguous genomic sequences, totaling 12,687 contigs of scaffold-scale lengths up to 1.1 Mb, and encompassing 783.4 Mb, or 98% of the genome length.

#### Pseudochromosome construction

To build chromosome length assemblies, the contigs from the final de novo assembly were anchored to an updated version of the *C. mollissima* reference genetic linkage map. We used the updated reference map (5) as the chestnut genome reference. The updated reference map includes additional SNP markers (net increase of 166 markers) not previously scored on the reference populations (‘Vanuxem’ x ‘Nanking’ and ‘Mahogany’ x ‘Nanking’ crosses), thus updating the current reference map (9).

To this updated reference map we attempted to add markers from three other *Castanea* linkage maps (three inter-specific chestnut maps from the crosses AD98 x KY115, CG61 x NCDOT, and ‘Cranberry’ x JB197), plus one high density oak genetic linkage map (6). The additional three chestnut maps were generated as part of a QTL mapping project for *Phytophthora cinnamomi* resistance (7). Linkage group orientation and marker order for these maps are in agreement with that of the updated reference map.

Our script for integrating these maps into the updated reference map included the following steps: (1) check all maps’ markers for LG assignment and linkage group orientation versus the updated reference map; (2) eliminate markers not assigned to the correct LG and reorient inverted LGs to match reference map orientations; (3) using markers in common (based on BLASTn search) between the individual maps and the reference map, fit each map to the updated reference map using transformation regression (Proc Transreg, SAS Institute, Cary, NC USA); (4) for each map fit, iteratively eliminate outlier markers and re-fit the models until no additional outliers are present, this resulted in new maps scaled to the same cM scale as the updated reference map; (5) for each scaffold containing one or more marker from the respective linkage maps scaled to the updated reference map, calculate the average updated reference map position; and (6) sort the scaffolds by scaled updated reference map position, providing the pseudochromosome build for each LG.

The updated reference map contained 1,322 markers, while the scaled-integrated map consisted of 4,283 markers. Marker sequences were located on contigs by BLASTN. Contigs were anchored to their chromosomal position based on the markers in the scaled-integrated map. If contigs were anchored to more than one chromosomal location, the best location was identified as follows: priority was given to the location indicated by a marker from the expanded *C. mollissima* reference map, followed by a marker from an interspecific map, followed by a marker from the *Q. robur* map. If the two possible locations were indicated by markers from the same map or markers on different interspecific maps, the marker with the best BLAST hit e-value to the sequence contig was selected for the final location.

### A.3. Chestnut Cytogenetics

#### Plant Materials and Chromosome Preparation

Actively growing root tips were excised from Chinese chestnut seedlings growing in potting soil. The excised root tips were pretreated with an aqueous solution of a-monobromonaphthalene (0.8 % v/v) and/or 2.5 mM 8-hydroxyquinoline for 1.5 h or 3 h, respectively, in the dark to accumulate prophase and metaphase stages for FISH, and then fixed in 4:1 (95% ethanol: glacial acetic acid). Fixed root tips were treated with cell wall degrading enzymes (40% (v/v) Cellulase (C2730, Sigma), 20% (v/v) Pectinase (P2611, Sigma), 40% (v/v) 0.01 M citrate buffer, pH 4.5, 2% (w/v) Cellulase RS (Yakult Pharmaceutical, Tokyo, Japan), 1% (w/v) Macerozyme (Yakult Pharmaceutical) and 1.5% (w/v) Pectolyase Y23 (Kyowa Chemical Co., Osaka, Japan)) and the chromosome spreads were prepared as described previously (8). For meiocyte pachytene chromosome spreads, emerging Chinese chestnut (genotype ‘Vanuxem’, ramet AD274) male flower buds (catkins) were harvested and placed in 3:1 ethanol:glacial acetic acid fixative and then transferred to fresh 3:1 fixative containing 1% Polyvinyl Pyrrolidone (BP431-100, Fisher Scientific, USA). Emerging anthers from the flower buds were isolated and squashed under a 22 × 22 mm glass cover-slip to provide the best anthers for pachytene analysis, and then intermittently heated over an alcohol burner and tapped with a forceps to spread the chromosomes. Slides with good chromosome spreads were stored at −80C for future use.

#### Probe Labeling and FISH

Probe DNAs (BAC clones from the Chinese chestnut physical map (2), Supplementary Table S3) including 18S-28S rDNA and 5S rDNA probes were labeled with biotin-16-dUTP (Biotin Nick Translation Mix, Roche, USA) and/or digoxigenin-11-dUTP (Dig Nick Translation Mix, Roche, USA) following the manufacturer’s instructions.

#### Fluorescence in situ Hybridization (FISH)

With BAC clones on somatic metaphase of different cultivars of Chinese chestnut and pachytene chromosome spreads of the indicated Chinese chestnut genotype was carried out as described elsewhere (9,10,8). Probe hybridization sites were detected with Cy3-conjugated streptavidin (Jackson ImmunoResearch Laboratories, USA) for biotin labeled probes and FITC-conjugated anti-digoxygenin (Roche, USA) for digoxigenin labeled probes. The FISH preparations were mounted with Vectashield containing DAPI (Vector Laboratories, USA) to prevent photo-bleaching the fluorochromes. Digital images were recorded using an epi-fluorescence microscope (AxioImager M2, Carl Zeiss, Germany) with suitable filter sets (Chroma Technology, USA) and a Cool Cube high performance CCD camera, and processed with ISIS V5.1 (MetaSystem Inc., USA) and Adobe Photoshop CS v8 (Adobe System, USA).

### A.4. Chinese chestnut genome functional features methods

#### Repeat masking and repetitive sequence annotations

Repeat masking was performed by first running RepeatModeler v1.0.10 using ‘ncbi’ as the engine to identify novel repeats (11). The RepeatModeler output was modified using a custom python script (12) to remove ribosomal RNAs (rRNAs). These novel repeats were concatenated with the RepBase plant repeat library (13), and input to RepeatMasker v4.0.6 with the parameters to ignore low complexity regions (-nolow) and to softmask repeats from the contigs (-xsmall) (14). The ‘ProcessRepeats’ software that comes with RepeatMasker was used to correctly classify the repeats from the RepBase plant library with the flag -species eudicotyledons.

#### RNA sequencing methods

Total RNA was extracted from leaf, petiole, twig, bark, and root tissues of the Vanuxem Chinese chestnut reference cultivar, from twig and leaf samples of the Nanking Chinese chestnut cultivar, and from roots of 25 cultivar Nanking seedlings which had been challenged with the *Phytophthora cinnamomi* root-rot pathogen, using the Qiagen RNeasy protocol (15). Total RNA samples were converted to HiSeq Illumina cDNA libraries using the TruSeq® Stranded Total RNA Library Prep kit for Plants. Quality checks of RNA and library preparations were conducted by micro-capillary electrophoresis on the 2100 BioAnalyzer (Agilent Genomics).

BioAnalyzer RIN value of 8.0 was used as minimum quality scores. The RNASeq libraries were selected for 250 bp insert sizes, and pooled prior to multiplex sequencing on a Illumina HiSeq 2000, producing 150 bp paired-end reads. Sequencing was conducted in rapid mode at the Pennsylvania State University Genomics Core Facility. Reads were demultiplexed into separate forward and reverse read FASTQ files using the barcoded adaptor sequences. The RNAseq reads were trimmed of adaptors and further filtered for base quality using the CLC Genomics Workbench (Qiagen) tools, selecting reads with a minimum quality score of 0.01 and minimum length of 100 bp. Two to three Gb of high quality RNAseq data was produced in each direction for each of the libraries. The filtered reads were aligned to the Vanuxem reference genome to check for alignment quality and contaminants prior to use in gene model predictions and validations.

#### Gene Prediction

BRAKER2 was used to identify gene models in the contigs (16). It6 was run with the soft-masked version of the contigs using two lines of evidence for training: alignments of C. mollissima RNASeq reads and alignments of the protein sequences from the Q. robur genome version PM1N, selected because it is the closest fully sequenced reference genome with high quality gene models (17). The RNASeq reads were aligned with STAR (18), and the proteins were aligned to the chestnut contigs with GenomeThreader (19).

Gene model predictions were sorted into high quality and low quality references by two criteria. First, expression evidence was examined. Using the gff file from BRAKER and the RNASeq alignments from STAR, the number of reads per predicted gene was assessed using HTSeq (20). Next, homology to *Q. robur* proteins was determined. A reciprocal BLASTp analysis was run to identify chestnut genes with a likely ortholog in *Q. robur* (21). Any gene model with at least 100 RNASeq aligned reads and/or a reciprocal best hit to a Q. robur gene was retained as high quality; the rest were placed in the low quality category.

#### Quality Assessment and Functional Annotation

Benchmarking Universal Single-Copy Orthologs (BUSCO) analysis was used to compare the gene model to 1,440 common orthologs across all embryophytes (22). To predict function, genes were annotated by BLAST searches (21) to SWISS-PROT and TrEMBL protein databases (e-value < 1e-5) (23), InterProScan sequence searches with Gene Ontology results parameter (24), and ghostKOALA searches against the Kyoto Encyclopedia of Genes and Genomes (KEGG) (25)

#### Comparative genome structure analyses

Orthogroups were built by OrthoFinder v2.2.7 (26) using the chestnut proteins and proteins from Arabidopsis thaliana version TAIR10 (27), Prunus persica v2.1 (28), Populus trichocarpa v3.1 (29), Vitis vinifera v2.1 (30), and Quercus robur version PM1N (17). The first four were downloaded from Phytozome (31), and the latter from the Oak Genome Sequencing website (32). Orthogroups containing genes from the monolignol biosynthetic pathway were flagged if they contained the genes annotated from that pathway reported in Arabidopsis thaliana (33) and/or Populus trichocarpa (34). The genes from poplar were converted from genome annotation version 1.1 to genome annotation version 3.1 by searching for the older gene name in Phytozome, and if not found, BLAST against the new version 3.1 genes. The version 3.1 gene was only accepted for BLAST results with at least 95% identity. NBS LRR genes were identified by searching the Chestnut genes with the Pfam model NB-ARC (PF00931.22) with HMMER v3.2.1 (35).

Circos plots were built using circos version 0.69-6 (36). The circos map of contigs to the Kubisiak reference genetic map used the same BLAST results used to anchor the contigs to chromosomes. The circos map of contigs to the Quercus robur and Prunus persica genomes were built using orthologs. The orthologs were identified by orthogroups with a single chestnut member gene and a single other species member gene in order to exclude gene families. Further filtering of the orthologs was performed to ensure only linkages with at least two points of agreement were retained. This was done by examining each ortholog in chestnut against the closest upstream and downstream ortholog. If either the upstream or downstream ortholog did not match the target genome on the same chromosome within 10Mb, the ortholog was discarded.

### A.6. Genome diversity and evolution methods

#### Sequencing and quality control of raw reads

For each of the 98 trees sampled, genomic DNA was extracted from leaf samples and paired-end sequencing libraries with insert sizes between 100 and 700 bp (an average of 350 bp) were constructed according to the Illumina library preparation protocol. We re-sequenced the individually indexed genomic libraries of all 98 trees to an expected depth of 25× per tree using the Illumina Genome Analyzer (HiSeq 2500), and a read length of 150 bp (paired-end). This generated a total of 13.4 billion 150 bp reads and an average of 136.4 million 150 bp reads (25.57× genome coverage) per tree.

The generated raw reads were subject to quality control using FASTX-toolkit (http://hannonlab.cshl.edu/fastx_toolkit/). Bases with Phred quality score <= 20 were defined as low quality. Low quality bases were masked, further were trimmed if were at the read end. Reads with low-quality bases > 95% of read length or with a length less than 30 bp were discarded and were also trimmed of any adaptor and repetitive telomere sequences.

#### Alignment and genotyping

High quality clean reads were mapped to the *C. mollissima* reference genome (https://www.hardwoodgenomics.org/chinese-chestnut-genome) using BWA-MEM (0.7.16a-r1181) with default setting (37), and alignment results were sorted and marked duplicate reads were removed using the Picard-tools v 1.92 (picard.sourceforge.net/). The mapping ratios for *C. mollissima*, *C. seguinii* and *C. henryi* were 95.2%, 96.1% and 96.7%, respectively.

The Genome Analysis Toolkit v 4.0.2.1 (GATK4) (38) was used to perform genotype calls individually. First, we used HaplotypeCaller command to call genotypes per site and generated a GVCF format file for each tree. Second, we used GenotypeGVCFs and SelectVariants commands to obtain a list of potential SNPs for each tree. Third, we used a hard-filtering approach to filter raw SNPs for each tree. We determine the filtering rule “DP < 8.0 || DP > 60.0 || QD < 2.0 || MQRankSum < −12.5 || ReadPosRankSum < −8.0 || MQ < 40.0 || FS > 60.0 || SOR > 3.0” by calculating the distribution of each statistics and used VariantFiltration command to perform the hard-filtering. If one SNP site is near an indel with distance < 3 bps, it was marked as missing. Finally, for homozygous-reference calls, we further filtered by minimum and maximum depth (DP < 8.0 and DP > 60.0). Therefore, through these filtration steps, we generated confident SNPs and also invariant homozygous sites for each tree. The indels identified by HaplotypeCaller command or other sites not satisfying the filtering criteria were documented as missing. For each tree, the set of confident SNPs and invariant sites were used to reconstruct FASTA format files, which were then used in the following population genomic analyses.

#### Validation of SNPs quality

Four individuals of *C. mollissima* and three individuals of *C. henryi* were used to assess the quality of our SNPs. For each tree, two separate DNA samples were sequenced. Then these were processed in identical pipeline with the processing from quality control to genotyping blind to the fact that they were duplicates. Then we compared the genotypes in the two samples for each tree. The rate of discordant genotypes between the two duplicate samples was very low (0.66, 0.63, 0.67, 0.94, 0.70, 0.74, 0.75 ×10^−3^ per SNP) for each tree, reflecting an average error rate lower than 0.37 ×10^−3^ per SNP. It suggested that the data set used in the analyses is of high quality.

#### Population size history

We inferred demographic history using SMC++ (39) based on population samples for each of three *Catanea* species in China, respectively. For each individual sample, whole genome diploid consensus sequence was generated from the confident SNPs and invariant homozygous sites. The mutation rate was set to 5 × 10^−9^ substitutions per site per year (40), and 15 years per generation was assumed. Note, if we use a lower estimate of the mutation rate, all results from the SMC++ and fastsimcoal2 (below) analyses remain qualitatively same but the estimated divergence time and effective population sizes are larger.

#### Individual clustering

We used phylogenetic analysis of the concatenated datasets, genetic assignment analysis, and comparison of species phylogenetic tree models to infer the relationships among three *Castanea* species. After removing sites with missing bases, a total of 2,599,011 fourfold degenerate sites with 165,244 polymorphic sites were retained and used in clustering analyses. The individual phylogenetic tree was reconstructed using RAxML-NG v. 0.6.0 with ‘GTR+FO+G4m’ model and 200 replicates of bootstrap analyses (41). In genetic assignment analysis, we ran ADMIXTURE ver. 1.23 (42) with cross-validation for values of genetic clusters *K* from 1 to 10 to infer the individual ancestry proportions with default settings. The optimum number of clusters (*K*) was determined using the cross-validation errors provided by ADMIXTURE ver. 1.23 (42).

In the coalescent analysis, three species phylogenetic tree topology were modeled and compared using fastsimcoal2 (43,44). Model 1 represents a close relationship between *C. mollissima* and *C. seguinii*. Model 2 represents a close relationship between *C. mollissima* and *C. henryi*. Model 3 represents a close relationship between *C. seguinii* and *C. henryi*. For each pair of species, 2D joint site frequency spectra were constructed and used in calculating likelihoods based on 300,000 coalescent simulations under each model. The highest likelihood and parameters were estimated in 200 ECM cycles. The mutation rate was set to 5 × 10^−9^ substitutions per site per year (40), and 15 years per generation was assumed. The “C” parameter, that is the minimum size of entry of the observed and simulated SFS, was set to 5 and the Akaike Information Criterion (AIC) values were used to rank three models. Standard deviations were determined from estimates of different C parameter settings from 1 to 20.

#### Introgression analysis

Introgression, that is the transfer of small genomic regions from a donor population / species into a recipient population/ species, is characterized by smaller divergence time at the introgressed locus, relative to the time of divergence from the donor species. Unfortunately, it is still challenging to exactly estimate divergence time for single introgressed locus. In neutral model, the relationship between absolute divergence dXY and divergence time t between X and Y, is defined as E(dXY)= 2µt+4µ*N*ANC (45), where *N*ANC is the effective population size of their ancestor before splitting. Thus, genomic regions with lower dXY level are thought to involve into introgression. However, lower mutation rate can also generate signals of lower dXY level at some genomic regions, which causes false positive inference of introgression. To avoid this false positive, we introduce an outgroup (O) to generate a scaled statistic *tit* which is defined as *tit* = dXY/(dXO + dYO). After reducing the mutation rate µ, *tit* becomes (t+2*N*ANC)/(2T+4*N*T), where *T* is the divergence time between X (Y) and the outgroup (O) and *N*T is the effective population size of their ancestral species at time *T*. For an introgressed gene or genomic region with introgression time *ti*, given the species divergence time at *t* (so *t > ti*), its *tit* statistic would be smaller than expected tit statistic values under neutral divergence model without gene flow because of (t+2*N*ANC)/(2T+4*N*T) > (ti+2*N*ANC)/(2T+4*N*T).

In the present study, we calculated *tit* per locus and used these results to infer introgressed genes. The significance level was determined with coalescent simulations. To generate the distribution of *tit* under neutral model, we estimated population parameters under the isolation with migration model based on the SFS generated from fourfold degenerate sites using fastsimcoal2 (43, 44). When calculating likelihoods, we performed 300,000 coalescent simulations for each of 200 ECM cycles. The mutation rate was set to 5 × 10^−9^ substitutions per site per year (40), and 15 years per generation was assumed. The “C” parameter was set to 5. Then we used these population parameters to generate simulated samples which were used to compute the expected distribution of *tit*. Additionally, we also compare the AICs of isolation with migration model and isolation without migration model to test whether gene flow occurred among species during their divergence history.

Strong selection in the ancestral population could reduce overall absolute divergence by reducing *θ*_ANC_=4µ*N*_ANC_. An extreme scenario is that, *θ*ANC decrease to zero. Thus, a lower tit or dXY can be explained by strong selection in the ancestral population. Given that strong selection occurred in the ancestral species of *C. mollissima* and *C. seguinii*, and to avoid this effect, we scaled simulated tit by multiplying t/(t+2*N*_ANC_).

The lineage-specific conserved regions/genes in both of the *C. mollissima* and *C. seguinii* genomes can also be used to explain the lower tit or dXY. To decrease this type of false positive, we computed the nucleotide diversity (π) for each gene and used π to infer conserved genes specific to the lineage including *C. mollissima* and *C. seguinii*. To avoid the noises from heterogeneous mutation rate among genes, we used a modified version of statistic r proposed by Innan (46), that is, scaled(π)= π/(dXO + dYO). The significant level was determined by coalescent simulations.

#### Population genetic test of selection at putative introgressed genes

We used the method of a modified HKA test to identify genes with unusual nucleotide polymorphism (46). First, we generated 100,000 samples using fastsimcoal (44) under the neutral model for *C. mollissima* and *C. seguinii*. Then we computed scaled(π) for each sample and each species and used them to produce the expected null distribution of scaled(π). These expected distributions were used to determine the significance level of HKA tests for each gene.

#### QTL selection signal analysis methods

Sequences of five accessions representing each American and Chinese chestnut species (see row 2, Supplementary Table S8) were pooled and aligned against reference Chinese chestnut (Vanuxem) genome assembly v.3.2. The Chinese chestnut genotypes, ‘Mahogany’ and ‘Nanking’ used as a donor of resistance to *Pc* and *Cp* in back-cross breeding program by TACF, were included into re-sequencing project. Sorted and indexed bam files were generated for each linkage group and ‘unmapped ‘contigs individually. The ANGSD software version 0.920 (htslib: 1.6; build Dec 5 2018 12:04:29) was used to calculate allele frequency spectrum, obtain a maximum likelihood estimate of the unfolded site frequency spectrum (SFS) (47), estimate pairwise nucleotide diversity and test for neutrality using Tajima’s D coefficient (48), which compares the number of pairwise differences to the number of segregating sites (49). Population genetic statistics was estimated for sliding 5kb windows along each linkage group with a step size of 1 kb. A window of 5 kb was selected because as 5-kb was an average size of gene models in a gff3 file. Output summary tables generated for *C. mollissima* and *C. dentata* were used to export Tajima’s D statistics and to calculate integrative indices of nucleotide diversity, i.e. ratios of *Pi Cden*/*Pi Cmol* and *Pi Cmol*/*Pi Cden*. To establish windows for *C. mollissima* vs. *C. dentata* and *C. dentata* vs. *C. mollissima* comparisons, cutoffs for 0.1%, 1 % and 5 % of most extreme *Pi*/*P*i values for empirical distribution across 12 linkage were set up for declaring significant departure from neutrality for the *Pi Cmol*/*Pi Cden*. and *Pi Cden*/*Pi Cmol* ratios respectively. QTL intervals tarits of interest (bud emergence, resistance to *Pc* and *Cp*) were delineated using sequence-based markers associated with QTLs and local BLAST^®^ (https://www.ncbi.nlm.nih.gov/books/NBK279690/) against the *C. mollissima* v3.2 contigs. Nucleotide diversity ratios nucleotide diversity (*PiCmiol/PiCden*, *PiCden/PiCmol*) ratios were calculated in excel and sorted using ‘subset’ command in R. Result of the neutrality test and (Tajima D) and nucleotide diversity (*Pi*/*Pi* ratios) were plotted using *ggplot2* in R (50).

## SUPPLEMENTARY TABLES AND FIGURES

**Table S1.**
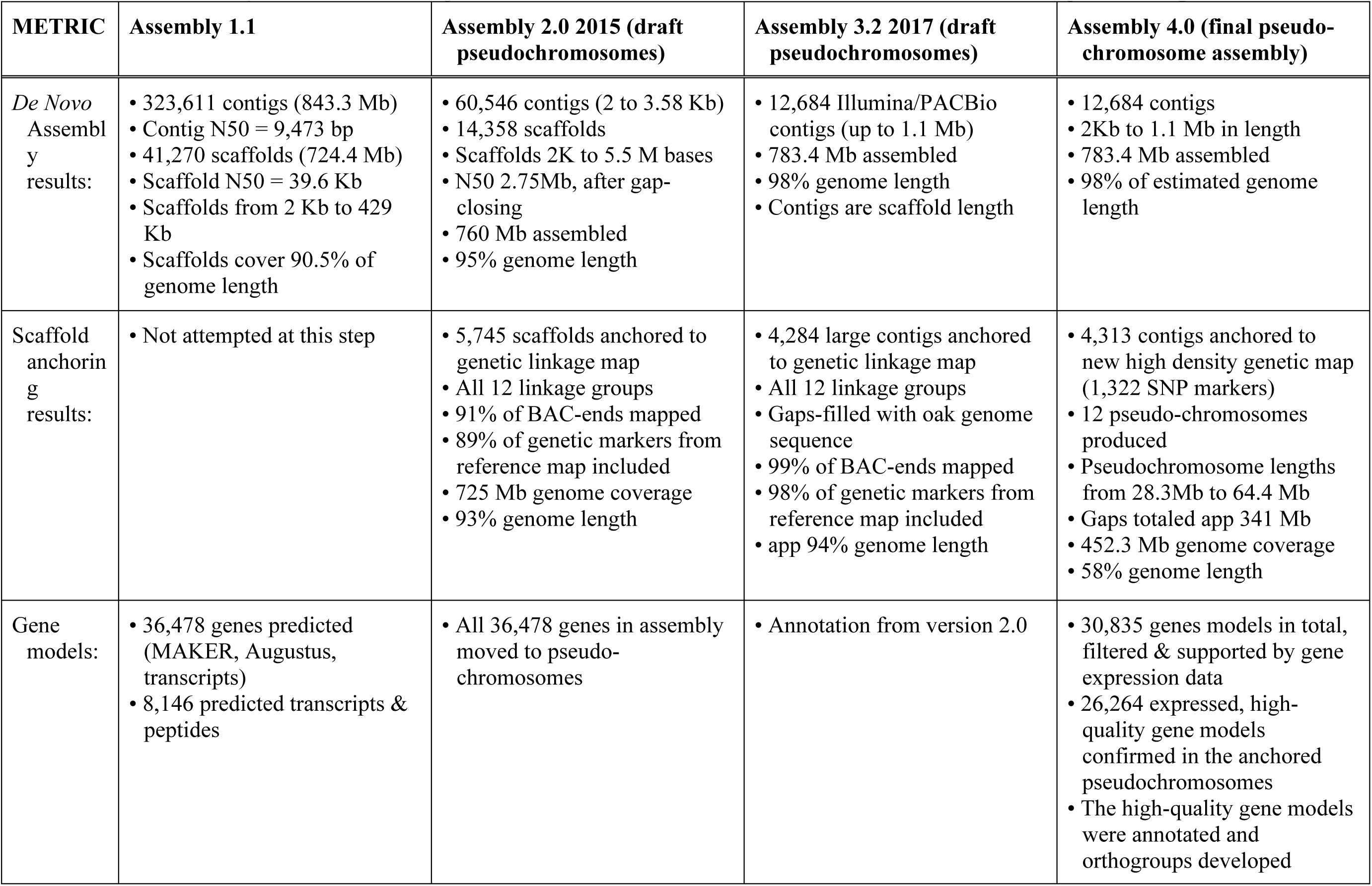
Metrics of key assemblies during construction of the C. mollissima cv. Vanuxem reference genome sequence.

| METRIC | Assembly 1.1 | Assembly 2.0 2015 (draft pseudo-chromosomes) | Assembly 3.2 2017 (draft pseudo-chromosomes) | Assembly 4.0 (final pseudo-chromosome assembly) |
| --- | --- | --- | --- | --- |
| *De Novo* Assembly results: | • 323,611 contigs (843.3 Mb)<br>• Contig N50 = 9,473 bp<br>• 41,270 scaffolds (724.4 Mb)<br>• Scaffold N50 = 39.6 Kb<br>• Scaffolds from 2 Kb to 429 Kb<br>• Scaffolds cover 90.5% of genome length | • 60,546 contigs (2 to 3.58 Kb)<br>• 14,358 scaffolds<br>• Scaffolds 2K to 5.5 M bases<br>• N50 2.75Mb, after gap-closing<br>• 760 Mb assembled<br>• 95% genome length | • 12,684 Illumina/PACBio contigs (up to 1.1 Mb)<br>• 783.4 Mb assembled<br>• 98% genome length<br>• Contigs are scaffold length | • 12,684 contigs<br>• 2Kb to 1.1 Mb in length<br>• 783.4 Mb assembled<br>• 98% of estimated genome length |
| Scaffold anchoring results: | • Not attempted at this step | • 5,745 scaffolds anchored to genetic linkage map<br>• All 12 linkage groups<br>• 91% of BAC-ends mapped<br>• 89% of genetic markers from reference map included<br>• 725 Mb genome coverage<br>• 93% genome length | • 4,284 large contigs anchored to genetic linkage map<br>• All 12 linkage groups<br>• Gaps-filled with oak genome sequence<br>• 99% of BAC-ends mapped<br>• 98% of genetic markers from reference map included<br>• app 94% genome length | • 4,313 contigs anchored to new high density genetic map (1,322 SNP markers)<br>• 12 pseudo-chromosomes produced<br>• Pseudo-chromosome lengths from 28.3Mb to 64.4 Mb<br>• Gaps totaled app 341 Mb<br>• 452.3 Mb genome coverage<br>• 58% genome length |
| Gene models: | • 36,478 genes predicted (MAKER, Augustus, transcripts)<br>• 8,146 predicted transcripts & peptides | • All 36,478 genes in assembly moved to pseudo-chromosomes | • Annotation from version 2.0 | • 30,835 genes models in total, filtered & supported by gene expression data<br>• 26,264 expressed, high-quality gene models confirmed in the anchored pseudo-chromosomes<br>• The high-quality gene models were annotated and orthogroups developed |

**Table S2.** Repetitive Regions of the Chinese Chestnut Assembly Version 3.2.

| Element Type | Number of Elements: | Length Occupied | Percent of Sequence |
| --- | --- | --- | --- |
| **Retroelements** | 169437 | 152637509 bp | 21.05% |
| SINEs: | 2504 | 241323 bp | 0.03 % |
| Penelope | 0 | 0 bp | 0.00 % |
| LINEs: | 43626 | 25668822 bp | 3.54 % |
| CRE/SLACS | 0 | 0 bp | 0.00 % |
| L2/CR1/Rex | 372 | 154244 bp | 0.02 % |
| R1/LOA/Jockey | 627 | 179751 bp | 0.02 % |
| R2/R4/NeSL | 0 | 0 bp | 0.00 % |
| RTE/Bov-B | 6541 | 1218084 bp | 0.17 % |
| L1/CIN4 | 35734 | 24071474 bp | 3.32 % |
| LTR elements: | 123307 | 126727364 bp | 17.48% |
| BEL/Pao | 1851 | 856235 bp | 0.12 % |
| Ty1/Copia | 59110 | 53059838 bp | 7.32 % |
| Gypsy/DIRS1 | 54496 | 66866289 bp | 9.22 % |
| Retroviral | 750 | 247176 bp | 0.03 % |
| **DNA transposons** | 129133 | 33416212 bp | 4.61 % |
| hobo-Activator | 60986 | 15501134 bp | 2.14 % |
| Tc1-IS630-Pogo | 0 | 0 bp | 0.00 % |
| En-Spm | 0 | 0 bp | 0.00 % |
| MuDR-IS905 | 0 | 0 bp | 0.00 % |
| PiggyBac | 0 | 0 bp | 0.00 % |
| Tourist/Harbinger | 11908 | 3930338 bp | 0.54 % |
| Other (Mirage, P-element, Transib) | 2511 | 307569 bp | 0.04 % |
| **Rolling-circles** | 0 | 0 bp | 0.00 % |
| **Unclassified:** | 711598 | 181937994 bp | 25.09% |
| **Total interspersed repeats:** | n/a | 367991715 bp | 50.74% |
| **Small RNA:** | 2620 | 376402 bp | 0.05 % |
| **Satellites:** | 724 | 124742 bp | 0.02 % |
| **Simple repeats:** | 3660 | 862105 bp | 0.12 % |
| **Low complexity:** | 0 | 0 bp | 0.00 % |

**Table S3.** LG specific BAC clones used in FISH to assign Chestnut chromosomes.

| **LGs** | **BAC clones** | **Map position** | **FISH BAC** | **Contig** |
| --- | --- | --- | --- | --- |
| A | BD162I11 | 0.55 cM | A-D12 | ctg 4835 |
|  | BD074J16 | 17.1 cM | A-C6 | ctg 645 |
|  | **Centromere Position: Tentative** |  |  |  |
|  | BB020L20 | 59.9 cM | A-A2 | ctg1683 |
|  | BD003A14 | 66.1 cM | A-C5 | ctg2116 |
|  | BB023C14 | 76.3 cM | A-C2 | ctg586 |
|  | BD157H16 | 80.70 cM | A-B12 | ctg3776 |
| B | BD110L01 | 6.15 cM | B-A10 | ctg3372 |
|  | BD075C12 | 9.4 cM | B-D6 | ctg3372 |
|  | BB033E09 | 13.2 cM | B-A3 | ctg2477 |
|  | **Centromere Position Tentative** |  |  |  |
|  | BB140H01 | 41.3 cM | B-A5 | ctg1982 |
|  | BB027F15 | 45.1 cM | B-E2 | ctg528 |
|  | BB166G01 | 63.75 cM | B-E6 | ctg12020 |
| C | BD017J05 | 11.60 cM | C-B7 | ctg4241 |
|  | BB033B24 | 11.60 cM | C-B2 | ctg4241 |
|  | **Centromere Position Tentative** |  |  |  |
|  | BD098P04 | 46.30 cM | C-E9 | ctg4032 |
|  | BD137D12 | 52.95 cM | C-A11 | ctg1820 |
| D | BD099P10 | 6.97 cM | D-F9 | ctg10919 |
|  | BD139F24 | 9.80 cM | D-C11 | ctg509 |
|  | **Centromere Position Tentative** |  |  |  |
|  | BB006G23 | 44.70 cM | D-C1 | ctg1251 |
|  | BD067M07 | 49.00 cM | D-H7 | ctg1212 |
| E | BD072B11 | 4.83 cM | E-A8 | ctg1231 |
|  | BB151N09 | 6.48 cM | E-C6 | ctg3296 |
|  | Centromere Position Tentative |  |  |  |
|  | BB078A16 | 51.00 cM | E-F3 | ctg4178 |
|  | BB124F09 | 53.70 cM | E-H4 | ctg10188 |
| F | BD157G22 | 0.90 cM | F-A12 | ctg1080 |
|  | BD136C19 | 7.53 cM | F-G10 | ctg2300 |
|  | Centromere Position Tentative |  |  |  |
|  | BD018C06 | 59.70 cM | F-C7 | ctg943 |
|  | BD154P01 | 59.70 cM | F-H11 | ctg943 |
| G | BD087O18 | 24.00 cM | G-B9 | ctg5364 |
|  | Centromere Position Tentative |  |  |  |
|  | BD073N19 | 56.40 cM | G-B8 | ctg5592 |
| H | BB134N22 | 6.33 cM | H-G6 | ctg4486 |
|  | BB171M04 | 6.33 cM | H-F10 | ctg4486 |
|  | Centromere Position Tentative |  |  |  |
|  | BD025A13 | 16.1 cM | H-G8 | ctg3854 |
|  | BB055C18 | 57.93 cM | H-F2 | ctg8575 |
| I | BD129K15 | 3.65 cM | I-E10 | ctg1454 |
|  | BD141N11 | 6.80 cM | I-D11 | ctg187 |
|  | Centromere Position Tentative |  |  |  |
|  | BD149F09 | 45.68 cM | I-G11 | ctg1727 |
| J | BB080D23 | 21.50 cM | J-G3 | ctg 285 |
|  | BD119H15 | 24.60 cM | J-C10 | ctg4883 |
|  | Centromere Position Tentative |  |  |  |
|  | BD085G21 | 56.9 cM | J-H8 | ctg2629 |
|  | BD083A05 | 58.3 cM | J-G8 | Ctg3140 |
| K | BB096M04 | 10.0 cM | K-C4 | ctg12025 |
|  | BD138E16 | 10.0 cM | K-B11 | ctg12025 |
|  | Centromere Position Tentative |  |  |  |
|  | BB025I23 | 59.60 cM | K-F1 | ctg2409 |
|  | BB062I20 | 60.00 cM | K-A3 | ctg563 |
| L | BD187B08 | 1.5 cM | L-H12 | ctg3391 |
|  | BD162A06 | 2.7 cM | L-C12 | ctg2087 |
|  | Centromere Position Tentative |  |  |  |
|  | BB072E18 | 47.0 cM | L-C3 | ctg4295 |
|  | BD077D03 | 47.0 cM | L-E8 | ctg4295 |

**Table S4.** Statistics of RNA-Seq Reads Aligned to Chinese Chestnut Assembly v 3.2.

| Sample Name | % Uniquely Mapped | % Multi Mapped | % Total Mapped |
| --- | --- | --- | --- |
| **Carlson (Penn State)** |  |  |  |
| Chestnut_Nanking_S15 | 69.21% | 2.51% | 71.72% |
| Chestnut_Vanuxem_S14 | 64.95% | 2.60% | 67.55% |
| **Sanger Sequencing** |  |  |  |
| CMFMEa | 97.18% | 1.41% | 98.59% |
| CMFMEb | 97.70% | 1.15% | 98.85% |
| CMFMEc | 98.06% | 0.87% | 98.93% |
| CMLMEa | 95.53% | 0.87% | 96.40% |
| CMLMEb | 94.91% | 1.44% | 96.35% |
| CMRMEa | 95.29% | 0.65% | 95.94% |
| CMRMEb | 94.68% | 0.75% | 95.43% |
| CM_Sanger_ESTs | 95.39% | 1.21% | 96.60% |
| CMSMEa | 94.43% | 1.32% | 95.75% |
| CMSMEb | 92.66% | 2.71% | 95.37% |
| **Holiday RNA-Seq** |  |  |  |
| JHC1 | 7.78% | 7.13% | 14.91% |
| JHC2 | 5.18% | 6.11% | 11.29% |
| JHC3 | 23.26% | 26.03% | 49.29% |
| JHC4 | 18.00% | 24.34% | 42.34% |
| JHC5 | 11.17% | 25.23% | 36.40% |
| JHC6 | 11.46% | 19.86% | 31.32% |
| JHC7 | 13.19% | 45.73% | 58.92% |
| JHC8 | 0.86% | 7.87% | 8.73% |
| JHC9 | 3.63% | 2.79% | 6.42% |
| **454 - *Castanea Mollissima*** |  |  |  |
| SRR006295 | 69.19% | 8.95% | 78.14% |
| SRR006296 | 67.33% | 23.48% | 90.81% |
| SRR006297 | 60.27% | 30.72% | 90.99% |
| SRR006298 | 55.49% | 25.65% | 81.14% |
| SRR006299 | 71.07% | 6.92% | 77.99% |
| SRR029309 | 58.35% | 29.93% | 88.28% |
| SRR029310 | 64.60% | 22.65% | 87.25% |
| **454 - *Castanea Dentata*** |  |  |  |
| SRR006300 | 75.52% | 10.49% | 86.01% |
| SRR006301 | 70.49% | 21.87% | 92.36% |
| SRR006302 | 71.97% | 21.79% | 93.76% |
| SRR006303 | 85.48% | 6.55% | 92.03% |
| SRR006304 | 85.76% | 6.45% | 92.21% |
| SRR006305 | 54.00% | 39.42% | 93.42% |
| SRR006306 | 78.15% | 13.70% | 91.85% |
| **Zhebentyayeva Root RNA-Seq** |  |  |  |
| tz10 | 82.82% | 3.03% | 85.85% |
| tz11 | 83.88% | 2.93% | 86.81% |
| tz1 | 84.13% | 2.69% | 86.82% |
| tz12 | 79.79% | 3.10% | 82.89% |
| tz13 | 83.71% | 2.77% | 86.48% |
| tz14 | 82.05% | 3.35% | 85.40% |
| tz15 | 79.94% | 4.14% | 84.08% |
| tz16 | 80.96% | 2.90% | 83.86% |
| tz17 | 82.88% | 3.39% | 86.27% |
| tz18 | 83.00% | 2.82% | 85.82% |
| tz2 | 82.96% | 2.78% | 85.74% |
| tz25 | 82.49% | 2.49% | 84.98% |
| tz26 | 81.22% | 3.14% | 84.36% |
| tz27 | 80.33% | 5.26% | 85.59% |
| tz28 | 81.24% | 2.68% | 83.92% |
| tz29 | 81.52% | 2.98% | 84.50% |
| tz30 | 80.81% | 2.81% | 83.62% |
| tz3 | 81.29% | 2.71% | 84.00% |
| tz4 | 86.17% | 3.63% | 89.80% |
| tz5 | 84.49% | 2.85% | 87.34% |
| tz6 | 84.28% | 3.14% | 87.42% |
| tz7 | 84.25% | 2.75% | 87.00% |
| tz8 | 82.90% | 3.34% | 86.24% |
| tz9 | 83.31% | 3.23% | 86.54% |

**Table S5.** OrthoFinder2 results of clustering orthologous groups of predicted proteins from the genomes of chestnut, Arabidopsis, peach, poplar, grape.

|  |  |
| --- | --- |
| Number of genes | 163,425 |
| Number of genes in orthogroups | 116,746 |
| Number of unassigned genes | 46,679 |
| Percentage of genes in orthogroups | 71.4% |
| Percentage of unassigned genes | 28.6% |
| Number of orthogroups | 16,687 |
| Number of species-specific orthogroups | 212 |
| Number of genes in species-specific orthogroups | 1,427 |
| Percentage of genes in species-specific orthogroups | 0.9% |
| Mean orthogroup size | 7.0 |
| Median orthogroup size | 6.0 |
| G50 (assigned genes) | 7 |
| G50 (all genes) | 6 |
| O50 (assigned genes) | 5077 |
| O50 (all genes) | 8805 |
| Number of orthogroups with all species present | 11,624 |
| Number of single-copy orthogroups | 3,122 |
| Number of orthogroups averaging <1 genes per-species | 2871 |
| Number of orthogroups averaging 1 genes per-species | 11428 |
| Number of orthogroups averaging 2 genes per-species | 1663 |
| Number of orthogroups averaging 3 genes per-species | 356 |
| Number of orthogroups averaging 4 genes per-species | 134 |
| Number of orthogroups averaging 5 genes per-species | 83 |
| Number of orthogroups averaging 6 genes per-species | 44 |
| Number of orthogroups averaging 7 genes per-species | 21 |
| Number of orthogroups averaging 8 genes per-species | 24 |
| Number of orthogroups averaging 9 genes per-species | 19 |
| Number of orthogroups averaging 10 genes per-species | 9 |
| Number of orthogroups averaging 11-15 genes per-species | 23 |
| Number of orthogroups averaging 16-20 genes per-species | 6 |
| Number of orthogroups averaging 21-50 genes per-species | 6 |

**Table S6.** Assignments of introgression-based candidate genes to Linkage Group B and cbr1 QTL contig positions by blast alignments. Two of the 7 candidate genes from the diversity analysis aligned to LGB, of which one gene (maker-scaffold00115-augustus-gene-0.47-mRNA-1) aligned to 6 places on LGB, outside of the cbr1 region, and the other (gene model snap_masked-scaffold01565-abinit-gene-0.16-mRNA-1), aligned to a single position with the major blight resistance QTL cbr1.

| Gene model | contig ID | Blast Eval | from | to | from | to | length (bp) |
| --- | --- | --- | --- | --- | --- | --- | --- |
| >maker-scaffold00115-augustus-gene-0.47-mRNA-1 | contig0002859 | 0 | 185 | 906 | 13567 | 12847 | This sequence maps to 6 loci (contigs) in LGB |
| >maker-scaffold00115-augustus-gene-0.47-mRNA-1 | contig0000499 | 3.22E-102 | 620 | 898 | 195789 | 195517 |  |
| >maker-scaffold00115-augustus-gene-0.47-mRNA-1 | contig0000176 | 1.51E-95 | 647 | 906 | 185259 | 185003 |  |
| >maker-scaffold00115-augustus-gene-0.47-mRNA-1 | contig0003568 | 5.43E-95 | 620 | 892 | 29738 | 30003 |  |
| >maker-scaffold00115-augustus-gene-0.47-mRNA-1 | contig0012709 | 6.08E-15 | 21 | 76 | 140941 | 140997 |  |
| >maker-scaffold00115-augustus-gene-0.47-mRNA-1 | contig0001071 | 7.86E-14 | 865 | 906 | 135795 | 135836 |  |
| >snap_masked-scaffold01565-abinit-gene-0.16-mRNA-1 | contig0001221 | 0 | 1 | 618 | 55909 | 55292 | in QTL locus |

\* Contigs identified using Kubisiak reference map (2013) markers were from *C. mollissima* Vanuxem v3.2 assembly
\*\* Contigs id'd using Fan markers were from *C. mollissima* (Vanuxem) v3.2 assembly
\*\*\* Contigs id'd using Sun gene models were from *C. mollissima* QTL assembly v1.1

**Table S7.**
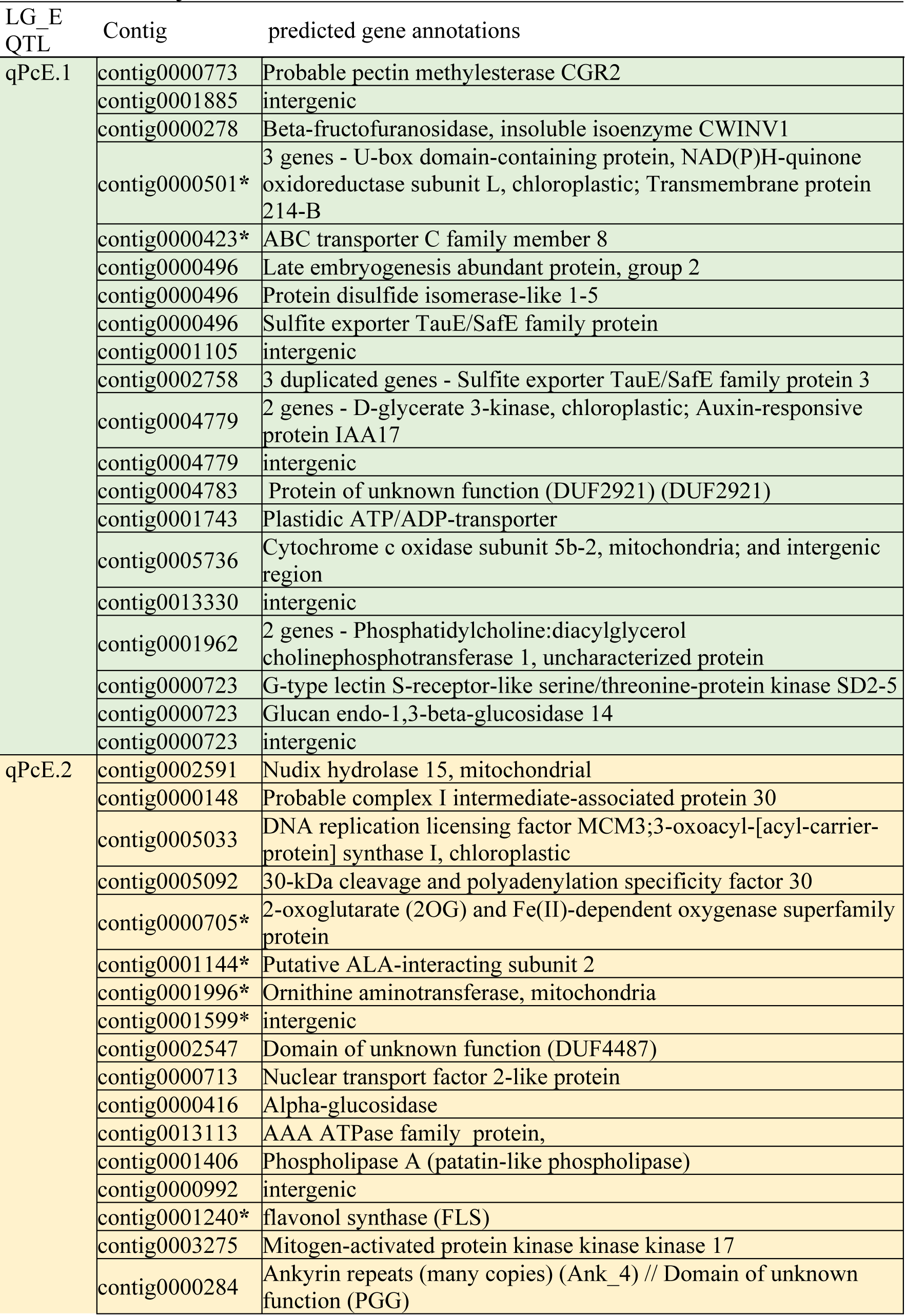
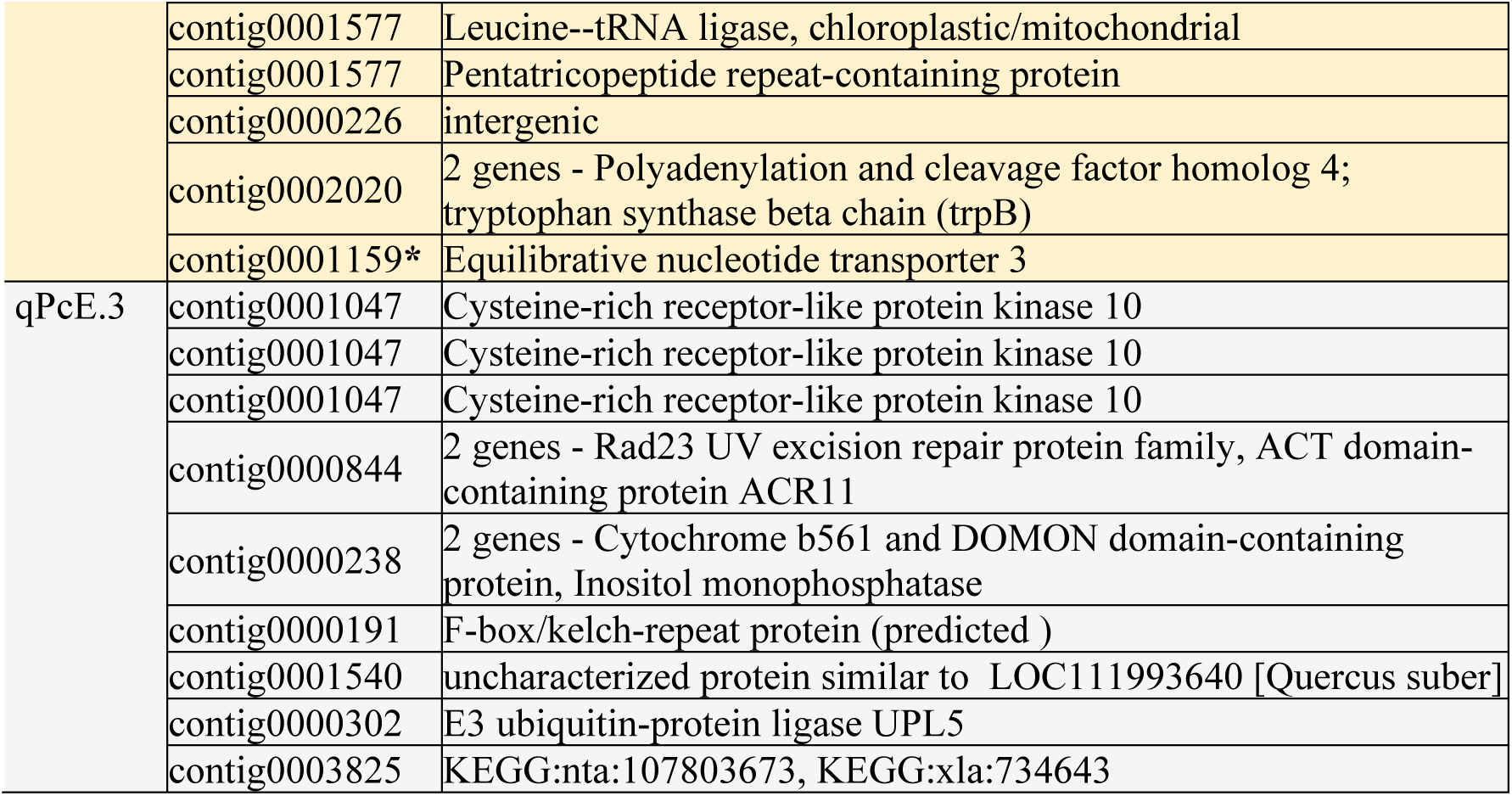
Selected candidate genes in C. *mollissima* Linkage Group E based on the distribution of Tajima D values relative to *C. dentata*.

| LG_E<br>QTL | Contig | predicted gene annotations |
| --- | --- | --- |
| qPcE.1 | contig0000773 | Probable pectin methylesterase CGR2 |
|  | contig0001885 | intergenic |
|  | contig0000278 | Beta-fructofuranosidase, insoluble isoenzyme CWINV1 |
|  | contig0000501* | 3 genes - U-box domain-containing protein, NAD(P)H-quinone oxidoreductase subunit L, chloroplastic; Transmembrane protein 214-B |
|  | contig0000423* | ABC transporter C family member 8 |
|  | contig0000496 | Late embryogenesis abundant protein, group 2 |
|  | contig0000496 | Protein disulfide isomerase-like 1-5 |
|  | contig0000496 | Sulfite exporter TauE/SafE family protein |
|  | contig0001105 | intergenic |
|  | contig0002758 | 3 duplicated genes - Sulfite exporter TauE/SafE family protein 3 |
|  | contig0004779 | 2 genes - D-glycerate 3-kinase, chloroplastic; Auxin-responsive protein IAA17 |
|  | contig0004779 | intergenic |
|  | contig0004783 | Protein of unknown function (DUF2921) (DUF2921) |
|  | contig0001743 | Plastidic ATP/ADP-transporter |
|  | contig0005736 | Cytochrome c oxidase subunit 5b-2, mitochondria; and intergenic region |
|  | contig0013330 | intergenic |
|  | contig0001962 | 2 genes - Phosphatidylcholine:diacylglycerol cholinephosphotransferase 1, uncharacterized protein |
|  | contig0000723 | G-type lectin S-receptor-like serine/threonine-protein kinase SD2-5 |
|  | contig0000723 | Glucan endo-1,3-beta-glucosidase 14 |
|  | contig0000723 | intergenic |
| qPcE.2 | contig0002591 | Nudix hydrolase 15, mitochondrial |
|  | contig0000148 | Probable complex I intermediate-associated protein 30 |
|  | contig0005033 | DNA replication licensing factor MCM3;3-oxoacyl-[acyl-carrier-protein] synthase I, chloroplastic |
|  | contig0005092 | 30-kDa cleavage and polyadenylation specificity factor 30 |
|  | contig0000705* | 2-oxoglutarate (2OG) and Fe(II)-dependent oxygenase superfamily protein |
|  | contig0001144* | Putative ALA-interacting subunit 2 |
|  | contig0001996* | Ornithine aminotransferase, mitochondria |
|  | contig0001599* | intergenic |
|  | contig0002547 | Domain of unknown function (DUF4487) |
|  | contig0000713 | Nuclear transport factor 2-like protein |
|  | contig0000416 | Alpha-glucosidase |
|  | contig0013113 | AAA ATPase family protein, |
|  | contig0001406 | Phospholipase A (patatin-like phospholipase) |
|  | contig0000992 | intergenic |
|  | contig0001240* | flavonol synthase (FLS) |
|  | contig0003275 | Mitogen-activated protein kinase kinase kinase 17 |
|  | contig0000284 | Ankyrin repeats (many copies) (Ank_4) // Domain of unknown function (PGG) |
|  | contig0001577 | Leucine--tRNA ligase, chloroplastic/mitochondrial |
|  | contig0001577 | Pentatricopeptide repeat-containing protein |
|  | contig0000226 | intergenic |
|  | contig0002020 | 2 genes - Polyadenylation and cleavage factor homolog 4; tryptophan synthase beta chain (trpB) |
|  | contig0001159* | Equilibrative nucleotide transporter 3 |
| qPcE.3 | contig0001047 | Cysteine-rich receptor-like protein kinase 10 |
|  | contig0001047 | Cysteine-rich receptor-like protein kinase 10 |
|  | contig0001047 | Cysteine-rich receptor-like protein kinase 10 |
|  | contig0000844 | 2 genes - Rad23 UV excision repair protein family, ACT domain-containing protein ACR11 |
|  | contig0000238 | 2 genes - Cytochrome b561 and DOMON domain-containing protein, Inositol monophosphatase |
|  | contig0000191 | F-box/kelch-repeat protein (predicted ) |
|  | contig0001540 | uncharacterized protein similar to LOC111993640 [Quercus suber] |
|  | contig0000302 | E3 ubiquitin-protein ligase UPL5 |
|  | contig0003825 | KEGG:nta:107803673, KEGG:xla:734643 |

**Table S8.** Tree materials utilized in the genome sequencing and subsequent analyses.

| **Objective** | **Tree material** |
| --- | --- |
| **1. Whole genome sequence reference tree** | ***C. mollissima* genotype ‘Vanuxem’ TACF** |
| **2. American and Chinese chestnut diversity panel for resequencing analyses of QTL regions for traits of importance** | ***C. dentata*: 5 trees** 03denGMBCLEM; 04denTFACLEM; 05denALRPENN; 06denHROPENN; 07denELLSUNY |
|  | ***C. mollissima*: 5 trees** 14molMHGCAES; 15molNKG TACF; 23molSPPCHNA; 27molFATPENN; 29molSTVPENN; 28molGILPENN |
| **3. Genome diversity and evolution of Asian chestnut species** | ***C. mollissima*: 43 trees/34 locations** |
|  | ***C. seguinii*: 28 trees/23 locations** |
|  | ***C. henryi*: 27 trees/19 locations** |

**Figure S1.**
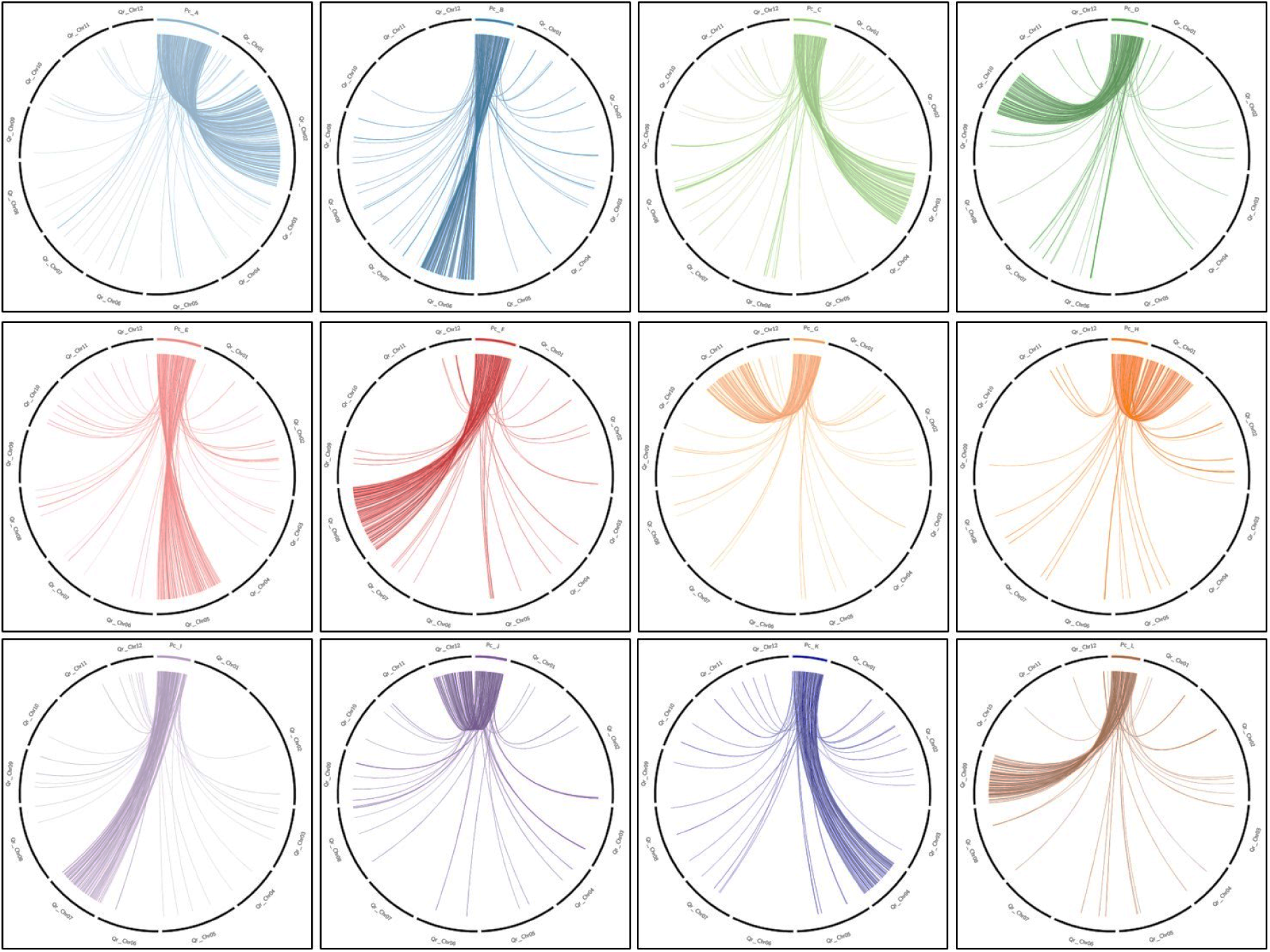
Circos plots of alignments of each C. mollissima pseudochromosome (Pc_A-L) vs all *Quercus robur* chromosomes (Qr_1-12). Pc, pseudochromosome. *C. mollissima* pseudochromosome naming convention of adheres to genetic linkage group (13) assignments.

**Figure S2.**
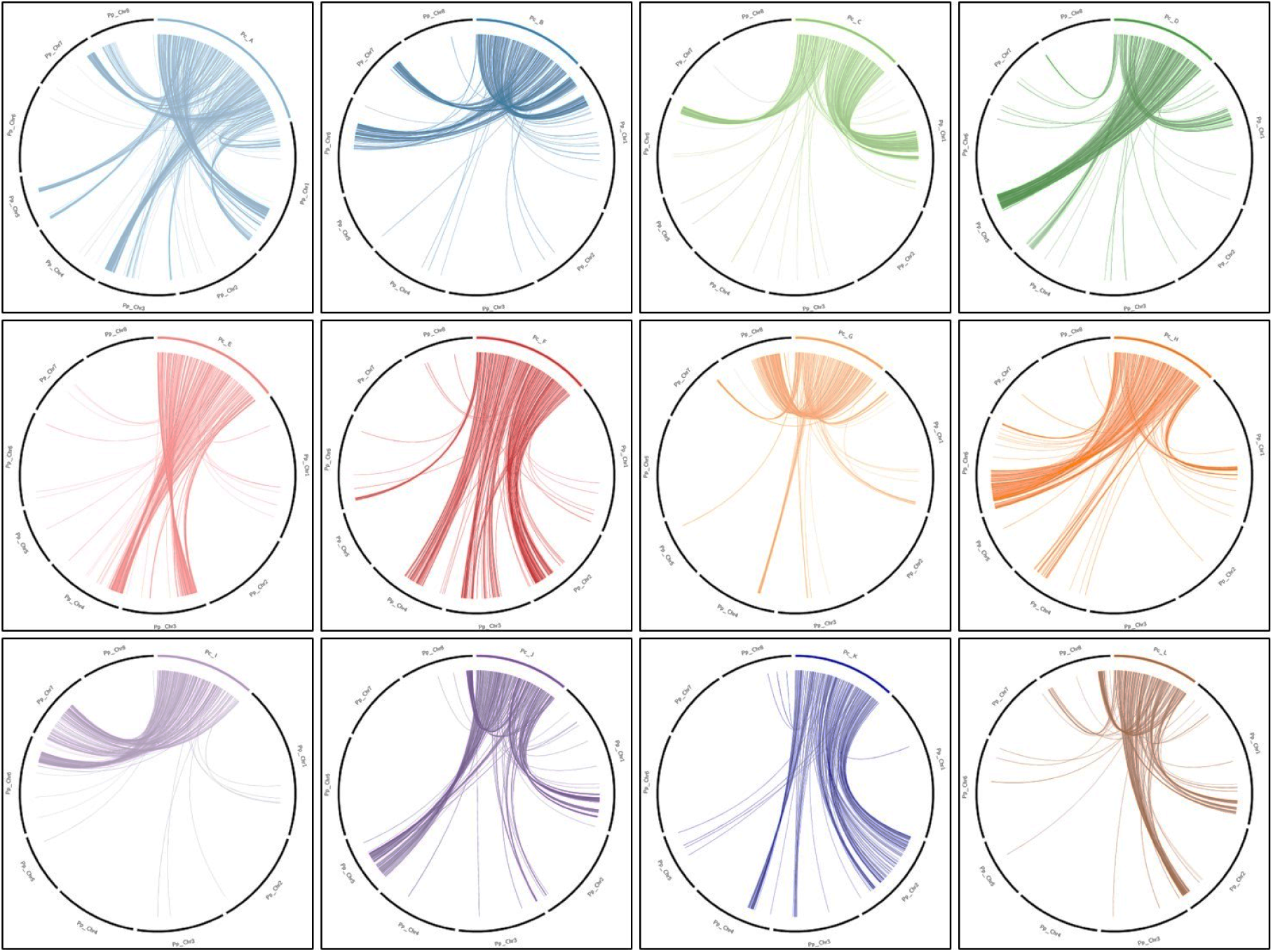
Circos plots of alignments of each *C. mollissima* pseudochromosome (Pc_A-L) vs all *Prunus persica* chromosomes (Pp_1-8). Pc, pseudochromosome. *C. mollissima* pseudochromosome naming convention of adheres to genetic linkage group (13) assignments.

